# Novel effector recognition capacity engineered into a paired NLR complex

**DOI:** 10.1101/2021.09.06.459143

**Authors:** Shanshan Wang, Weijie Huang, Zane Duxbury, Saskia A. Hogenhout, Jonathan D. G. Jones

**Affiliations:** The Sainsbury Laboratory, University of East Anglia, Norwich Research Park, Norwich NR4 7UH, United Kingdom; Department of Crop Genetics, John Innes Center, NR4 7UH Norwich, United Kingdom; Jealott’s Hill International Research Centre, Syngenta, Bracknell RG42 6EY, United Kingdom

**Keywords:** synthetic NLR, RRS1/RPS4, phytoplasma, SAP05, plant immunity

## Abstract

The Arabidopsis *RRS1-R Resistance* gene confers recognition of the bacterial acetyltransferase PopP2 and another bacterial effector, AvrRps4. The *RRS1-S* allele recognizes AvrRps4 but not PopP2. *RRS1- R/RRS1-S* heterozygotes cannot recognize PopP2. *RRS1-R* and *RRS1-S* also suppress the constitutive RPS4-dependent autoactivity of *RRS1-R*^*slh1*^. Phytoplasmas cause important plant diseases, and their effectors can cause degradation of specific host proteins. We tested whether attaching a pathogen effector-dependent degron to RRS1-R, enabling its degradation by phytoplasma effector SAP05, could derepress RRS1-R^*slh1*^ autoactivity, resulting in SAP05-dependent resistance. In transient assays in tobacco, RRS1-R-derived constructs can confer a hypersensitive response (HR) to SAP05. However, phytoplasma infection assays in transgenic Arabidopsis resulted in delayed disease symptoms but not full resistance. We provide a proof-of-concept strategy utilizing the recessiveness of a plant immune receptor gene to engineer recognition of a pathogen effector that promotes degradation of a specific host protein.

## Introduction

Plant immunity to pathogens involves both cell surface and intracellular receptors^1^. The cell surface receptors perceive (mostly) conserved pathogen-associated molecular patterns (PAMPs), and intracellular receptors recognise pathogen virulence factor (“effector”) proteins delivered into the cell^1^. In both plants and animals, the intracellular nucleotide-binding leucine-rich repeat receptor (NLR) family of proteins contributes to innate immunity^2^. Plant NLRs are activated upon perception of effector proteins. Activated NLRs lead to various defense responses towards pathogens, including a hypersensitive cell death response (HR) that restricts pathogens to the infection site, and the activation of transcription of defense-related genes. Since the isolation of the NLR-encoding resistance genes *N* from *Nicotiana tabacum*^3^, and *RPS2* from *Arabidopsis thaliana*^4^, hundreds of plant NLR genes have been cloned^5^. However, unlike most cell surface receptors, NLRs recognise specific cognate effectors from recognized pathogen races. Expanding the recognition spectrum of NLRs might provide novel resistance resources for future crop disease prevention.

There have been important recent advances in our understanding of how NLRs recognise their cognate effectors and activate immunity. Most plant NLRs carry an N-terminal Toll/interleukin-1 receptor/resistance protein (TIR) domain, a coiled coil (CC) domain or an RPW8 domain. They share a central nucleotide-binding site (NBS), which switches the NLR from inactive to active state (often correlated with ATP binding), a C-terminal leucine-rich repeat (LRR) domain and other C-terminal domains for effector recognition and self-regulation of the NLR structure. Some NLRs carry an integrated decoy domain that mimics authentic virulence targets of pathogen effectors^6-9^.

The structures of activated forms of NLRC4, a mammalian NLR, and plant NLRs ZAR1, Roq1 and RPP1 demonstrate that upon effector recognition, NLR proteins undergo conformational changes and form oligomers which impose induced proximity on their N-terminal signalling domains, resulting in activation of downstream immune signalling^9-13^. The conserved architectures and molecular mechanism of NLRs suggest the possibility of moving NLRs between species to confer new resistances^14^.

Wild species constitute a disease resistance (R) gene resource. However, time and effort are required to clone new *R* genes, and to validate the function and dissect the molecular mechanism of each NLR. Therefore, engineering plant NLRs is an attractive alternative potential route to gain novel resistance.

Gain of function mutagenesis screens have been performed to select alleles of Rx, R3a, I2 or L6 conferring response to new effectors^15-18^. Arabidopsis RPS5 monitors the guardee PBS1 for cleavage by AvrPphB from *Pseudomonas syringae*^19^, and modification of the PBS1 proteolytic target site to other proteolytic target sites enables immune response to AvrRpt2 from *P. syringae*, tobacco etch virus (TEV) and turnip mosaic virus (TuMV)^20^. Pik and Pia, paired NLRs from rice, with a Heavy Metal Associated (HMA) domain integrated in the sensor NLR Pik, function to recognise effector from *Magnaporthe oryzae*^21^. Modification of the HMA domain expanded the range of effector recognition^22-25^.

In Arabidopsis, RRS1-R (*Resistance to Ralstonia solanacearum* 1) and RPS4 (*Resistance to Pseudomonas syringae* 4) are well-studied paired TIR-NLRs, functioning together to recognize bacterial effector AvrRps4 from *P. syringae*, PopP2 from *R. solanacearum* and an unknown effector from *Colletotrichum higginsianum*^26-28^. The effectors are recognized via direct interaction with and/or modification of the integrated C-terminal WRKY domain of RRS1, followed by defense activation through RPS4^29-32^. The Arabidopsis RRS1-R^*slh1*^ (sensitive to low humidity 1) mutant with a single leucine insertion in WRKY domain causes an RPS4-dependent effector-independent autoimmune phenotype^33,34^. Most NLR genes are dominant, but like RRS1-R, RRS1-R^*slh1*^ is also recessive, which means the autoimmune phenotype can be suppressed by inactive forms of RRS1 such as RRS1-R from Arabidopsis Ws-2 or RRS1-S from Col-0. The RRS1*-*R^*slh1*^-induced-RPS4-dependent autoactivity (hereafter RRS1*-*R^*slh1*^ autoactivity) overlaps strongly with AvrRps4- and PopP2-induced RRS1/RPS4 mediated response at transcriptional level^34^, and RRS1-R^*slh1*^ autoactivity could enable engineering effector-triggered immunity.

Here we designed a system consisting of an autoactive RRS1-R^*slh1*^, an RPS4 and an inactive RRS1 with an extra domain which can be recognised and degraded by the cognate effector. Upon effector action, RRS1-R^*slh1*^ autoimmunity is derepressed, and defense responses activated. To create this, we selected several effectors and their corresponding targets. SAP05, an effector from insect-transmitted phytopathogenic bacteria, degrades GATA zinc finger domain transcription factor proteins ^35^. We fused a GATA SAP05-dependent degron domain to RRS1-R. Co-expression of this fusion protein represses autoimmunity of RRS1-R^*slh1*^; this is derepressed by coexpression with SAP05, restoring the RRS1-R^*slh1*^ -induced-RPS4-dependent hypersensitive response (HR) in transient assays. Transgenic Arabidopsis with the engineered resistance proteins show delayed disease symptoms after phytoplasma infection though not full resistance. This synthetic NLR combination results in new effector recognition capacity, and this approach could be applied to recognise a broad range of effectors from phytopathogens and serve as new source of plant resistance.

## Results

### Constructing an RRS1-derived allele that represses the autoactivity of RRS1-R^*slh1*^

We set out to engineer a system that contains three components; an autoactive RRS1-R^*slh1*^, RPS4, and an RRS1-derived variant carrying an effector target domain, here designated as RRS1-X (Fig. 1a). Effector action promotes degradation of RRS1-X, thus derepressing the RRS1-R^*slh1*^ autoactivity. RRS1 and RPS4 are nuclear localized NLRs, so nuclear-localized host targets and effectors were prioritized for evaluation. Transcription factors (TFs) are often targeted by effectors, which can lead to their degradation (Table. S1). For example, the phytoplasma effector SAP11 interacts with and destabilizes multiple TEOSINTE BRANCHED 1-CYCLOIDEA-PROLIFERATING CELL FACTOR (TCP) transcription factors^36-38^, SAP54 directly binds and promotes degradation of MADS domain transcription factor (MTF) family proteins, including SEP3, AP1 or SOC1^39^, and SAP05 directly binds the zinc finger domains of SQUAMOSA promoter-binding proteins (SBP) and GATA domain transcription factors and mediates their degradation^35^. *Pseudomonas syringae* Type III Effector HopBB1 promotes the degradation of TCP14 by direct binding^40^, and HopX1 eliminates JAZ proteins by cleaving the ZIM domain^41^.

**Figure 1.**
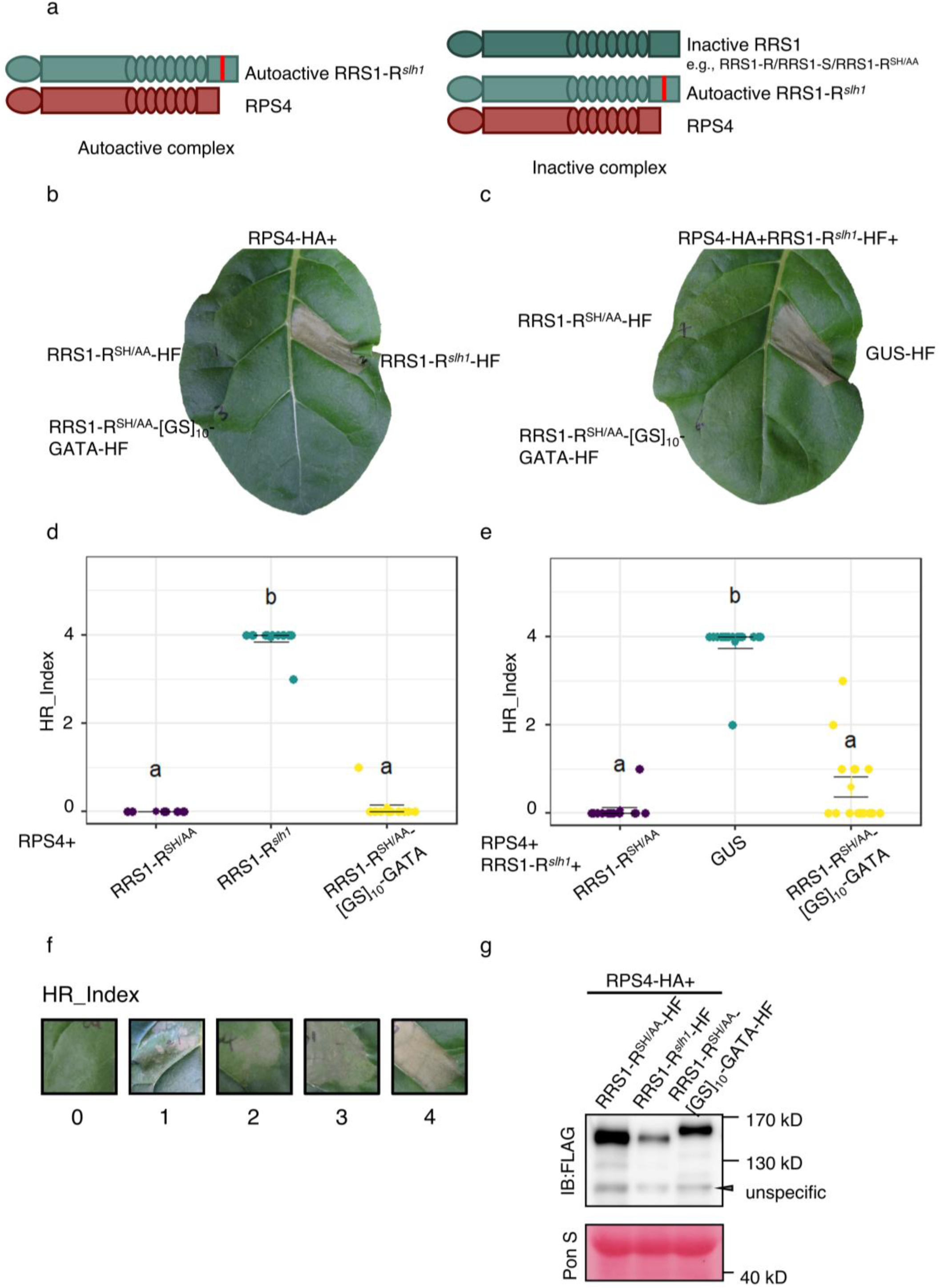
RRS1-R^SH/AA^-[GS]_10_-GATA suppresses RRS1-R^*slh1*^ autoactivity. **(a)** Schematic diagram of RRS1/RPS4 complexes. RPS4-dependent autoactivity of RRS1 (RRS1-R^*slh1*^) can be suppressed by inactive RRS1 allele (RRS1-R/RRS1-S/RRS1-R^SH/AA^). **(b)** HR assay in *N. tabacum* shows RRS1-R^SH/AA^-[GS]_10_-GATA-HF is not autoactive. **(c)** HR assay in *N. tabacum* shows RRS1-R^SH/AA^-[GS]_10_-GATA-HF suppresses RPS4-dependent RRS1-R^*slh1*^ autoactivity. **(b** and **c)** *N. tabacum* leaves were coinfiltrated with A. tumefaciens strains carrying the indicated constructs. Photos were taken 3-4 days post infiltration (dpi). **(d** and **e)** HR index of visually assessed tobacco infiltrations for (**b**) and (**c**) respectively. Each sample represents co-infiltrations of agrobacteria with different construct (indicated on the x axis) with a sample size of n= 12, 12 and 12 for (**d**) and n=15, 15 and 15 for (**e**). Errorbars show SD. Difference of HR levels was assessed by One way ANOVA (Analysis of variance) followed by a Turkey HSD test. Different letters indicate significant difference at level *p* < 0.05. **(f)** Representative image of HR index used to quantify cell death in tobacco leaf. **(g)** Protein expression analysis of the constructs. *N. benthamiana* leaves were coinfiltrated with A. tumefaciens strains carrying the indicated constructs and harvested after 48 hours post infiltration. Total proteins were immunoblotted with anti-FLAG antibodies for RRS1-R^SH/AA^-HF (His-FLAG), RRS-R^*slh1*^-HF or RRS1-R^SH/AA^-[GS]_10_-GATA-HF.

The C-terminus of RRS1 plays a crucial role in maintaining the inactive state of the RRS1/RPS4 complex as well as in responses to effectors^31,32^. The RRS1-R allele has an additional 83 amino acids at the C terminus compared to RRS1-S. Both RRS1-R and RRS1-S repress the autoactivity of RRS1-R^*slh1*^. Therefore, RRS1-R or RRS1-S were fused with target proteins/domains. Since NLR protein derivatives can easily lose function or become autoactive, a variety of linkers including Glycine Serine [GS] tandem repeat sequences, Asn-Ala-Ala-Ile-Arg-Ser [NAAIRS] tandem repeat sequences^42,43^ or a combination of both were applied to link RRS1 to the target protein/domain.

We transiently co-expressed RRS1 fusion proteins with RPS4 using agrobacterial infiltration into *Nicotiana tabacum* leaves to test for RRS1 fusion protein autoactivity. RRS1-R-[GS]_10_-TCP14, RRS1-R-[GS]_10_-JAZ1 and RRS1-R-[GS]_10_-GFP show RPS4-dependent autoactivity (Fig. S1a). We hypothesized this may be due to the size of the fused protein JAZ1 (253 amino acids), TCP14 (489 amino acids) and GFP (238 amino acids). The minimum size of target protein domain which responds to effector was then investigated. A keratin-like domain (KD) from MADS box transcription factors is sufficient to interact with phytoplasma effector SAP54^39^ and was fused to RRS1. Fusion proteins of RRS1-R carrying the KD with 3 different linkers show strong autoactivity when transiently co-expressed with RPS4 in *N. tabacum* (Fig. S1b) and the same strong RPS4-dependent autoactivity is observed with RRS1-S allele and KD fusion protein (Fig. S1c). We also fused the SBP domain from squamosa promoter-like 11 (SPL11) and GATA domain from GATA18, which are the targets of phytoplasma effector SAP05, to RRS1-R. All the tested fusion proteins show RPS4-dependent autoactivity, though RRS1-R-[GS]_10_-GATA-HF (His-FLAG) showed weak autoactivity (Fig. S1d). These results suggest that RRS1 structure is tightly self-regulated by intramolecular domain interactions and extra domains may disrupt the inactive state of RRS1/RPS4 complex.

### RRS1-R^SH/AA^ eliminates the autoactivity of the fusion protein

We tested an alternative method to eliminate the autoactivity of the RRS1 fusion protein. The structure of RRS1 and RPS4 TIR domains reveals that the heterodimerization of RPS4 TIR and RRS1 TIR is dependent on the integrity of the αA- and αE-helices (the AE interface), and mutations disrupting the interface prevent the signalling induction of RRS1/RPS4^44^. Because RRS1-R^SH/AA^, with H25A S26A mutations in the AE interface, abolishes RRS1/RPS4 function, but can repress RRS1-R^*slh1*^ autoactivity (Fig. 1a)^34,44^, we tested if these mutations eliminate the autoactivity of the fusion protein. Unlike the autoactive RRS1-R-[GS]_4_-[NAAIR]_2_-[GS]_4_-KD-HF, RRS1-R^SH/AA^-[GS]_4_-[NAAIR]_2_-[GS]_4_-KD does not show autoactivity with RPS4 (Fig. S2a). The TCP domain of transcription factor TCP2 (TCPd) or TCP18 (TCP18d), which is sufficient to associate with SAP11^36^, and MED19a which is destabilized by *Hyaloperonospora arabidopsidis* effector HaRxL44, were fused to RRS1-R^SH/AA^ respectively^36,45^. A 95 amino-acid unstructured region (Unstr) from *S. cerevisiae* cytochrome *b*_2_ was appended to the C terminus of TCPd to facilitate a rapid proteasome-dependent degradation^46^. RRS1-R^SH/AA^-[GS]_10_-TCP2d-FLAGUnstr, RRS1-R^SH/AA^-[GS]_10_-TCP18d-FLAG-Unstr, RRS1-R^SH/AA^-[GS]_10-_GATA-HF and RRS1-R^SH/AA^-[GS]_10_-MED19a were not autoactive when co-expressed with RPS4 in *N. tabacum* (Fig. S2a, Fig. 1b and 1d). Co-expression of RRS1-R^*slh1*^ and RPS4, RRS1-R^SH/AA^-[GS]_10_-GATA fully suppress RRS1-R^*slh1*^ autoactivity (Fig. 1c and 1e), whilst with the rest, only partial suppression was observed (Fig. S2b and S3a). Protein expression level were verified by western blot (Fig. 1g and S2c and S3b). Intriguingly, RRS1-R^SH/AA^-[NAAIR]_4_-GATA-HF cannot suppress RRS1-R^*slh1*^ autoimmunity (Fig. S3a), suggesting that the linker region is important to maintain the inactive state of the complex.

### Constructing a RRS1-R^SH/AA^ allele with a GATA domain that promotes SAP05-dependent degradation

To determine whether RRS1-R^SH/AA^-[GS]_10_-GATA-HF responds to effector SAP05, we performed transient expression assays in *N. tabacum* and found that expressing RRS1-R^SH/AA^-[GS]_10_-GATA-HF, RRS1-R^*slh1*^, RPS4 with SAP05 but not with control GUS-HF, triggered HR (Fig. 2a and 2b). RRS1-R^SH/AA^-HF did not respond to SAP05 (Fig. S4). Consistent with the recent report that SAP05 binds and subsequently degrades GATA domain-containing proteins ^35^, RRS1*-*R^SH/AA^-[GS]_10_-GATA-HF, but not RRS1*-*R^SH/AA^-HF, co-immunoprecipitated with SAP05 (Fig. 2c). RRS1*-*R^SH/AA^-[GS]_10_-GATA-HF, but not RRS1*-*R^SH/AA^-HF, was degraded when co-expressed in *N. benthamiana* with SAP05 in the presence of RPS4-HA and RRS1-R^*slh1*^-V5 (Fig. 2d). Transient assays in *N. benthamiana* and *N. tabacum* confirmed that in the presence of RPS4 and RRS1-R^*slh1*^, SAP05 mediates RRS1*-*R^SH/AA^-[GS]_10_-GATA degradation and therefore can derepress RRS1*-*R^*slh1*^ autoactivity.

**Figure 2.**
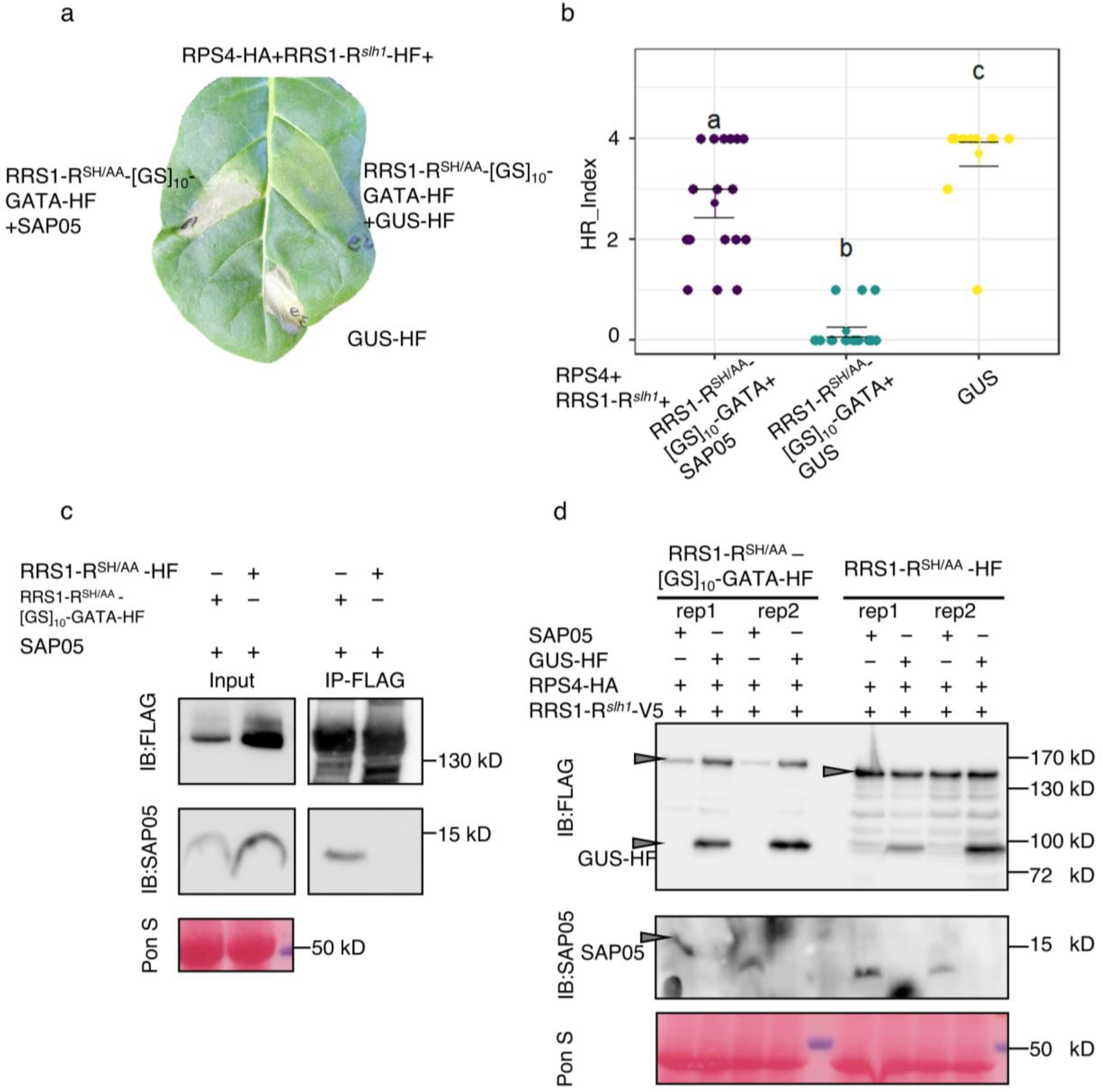
RRS1-R^SH/AA^-GATA responds to SAP05 and derepresses the RPS4-dependent RRS1-R^*slh1*^ autoactivity. **(a)** RRS1-R^*SH/AA*^-[GS]_10_-GATA restores RRS1-R^*slh1*^-mediated RPS4-dependent HR when coinfiltrated with SAP05. *N. tabacum* leaves were coinfiltrated with A. tumefaciens strains carrying the indicated constructs. Photos were taken at 4 dpi. **(b)** HR index of visually assessed tobacco infiltrations of (**a**). Each sample represents co-infiltrations of agrobacteria with different construct (indicated on the x axis) with a sample size of n= 17, 17 and 13. Errorbars show SD. Difference of HR levels was assessed by One way ANOVA (Analysis of variance) followed by a Turkey HSD test. Different letters indicate significant difference at level *p* < 0.05. (**c**) RRS1-R^*SH/AA*^-[GS]_10_-GATA interact with SAP05 in planta. *N. benthamiana* leaves were transiently coinfiltrated with A. tumefaciens strains carrying SAP05, RRS1-R^*SH/AA*^-[GS]_10_-GATA-HF or RRS1-R^*SH/AA*^-HF (as indicated by a + or -symbol). Two days after infiltration, the samples were harvested and proteins with FLAG epitope were immunoprecipitated and subjected to SDS-PAGE. Proteins were immunoblotted by anti-FLAG or anti-SAP05 antibody. Rubisco stained by Ponceau (Pon S) served as loading control. This experiment was performed twice with the same result. **(d)** RRS1-R^*SH/AA*^-[GS]_10_-GATA is degraded by SAP05 in planta. *N. benthamiana* leaves were transiently coinfiltrated with A. tumefaciens strains carrying RPS4-HA, RRS1-R^*slh1*^-V5, SAP05, GUS-HF, RRS1-R^*SH/AA*^-[GS]_10_-GATA-HF or RRS1-R^*SH/AA*^-HF (as indicated by a + or -symbol). Rubisco stained by Ponceau (Pon S) served as loading control. Samples were harvested at 3 dpi and the total protein were subjected to SDS-PAGE and immunoblotted by anti-FLAG or anti-SAP05 antibody. Two replicates were shown here. The experiment was repeated for three times with similar result.

### Transgenic Arabidopsis with RRS1-R^SH/AA^-[GS]_10_-GATA show delayed disease symptoms after phytoplasma infection

Using Golden Gate cloning^47^, binary vectors carrying RRS1*-*R^*slh1*^-V5 with RRS1*-*R^SH/AA^-[GS]_10_-GATA-HF (SH/AA-GATA) or RRS1*-*R^SH/AA^-HF (SH/AA) were generated for Arabidopsis transformation (Fig. 3a). We use the pAtSSR16 (AT4G34620) promoter to drive the RRS1 alleles and FastRed as selection marker^48^. The constructs were transformed into *Col-0/rrs1-3/rrs1b-1* background. Growth of the transgenic plants was indistinguishable from Col-0 or *Col-0/rrs1-3/rrs1b-1*(Fig. S5a). Two independent transgenic Arabidopsis lines were used for phytoplasma infection assays, of which 6 individual plants were sampled to confirm the protein expression by Western blot (Fig. S5b).

**Figure 3.**
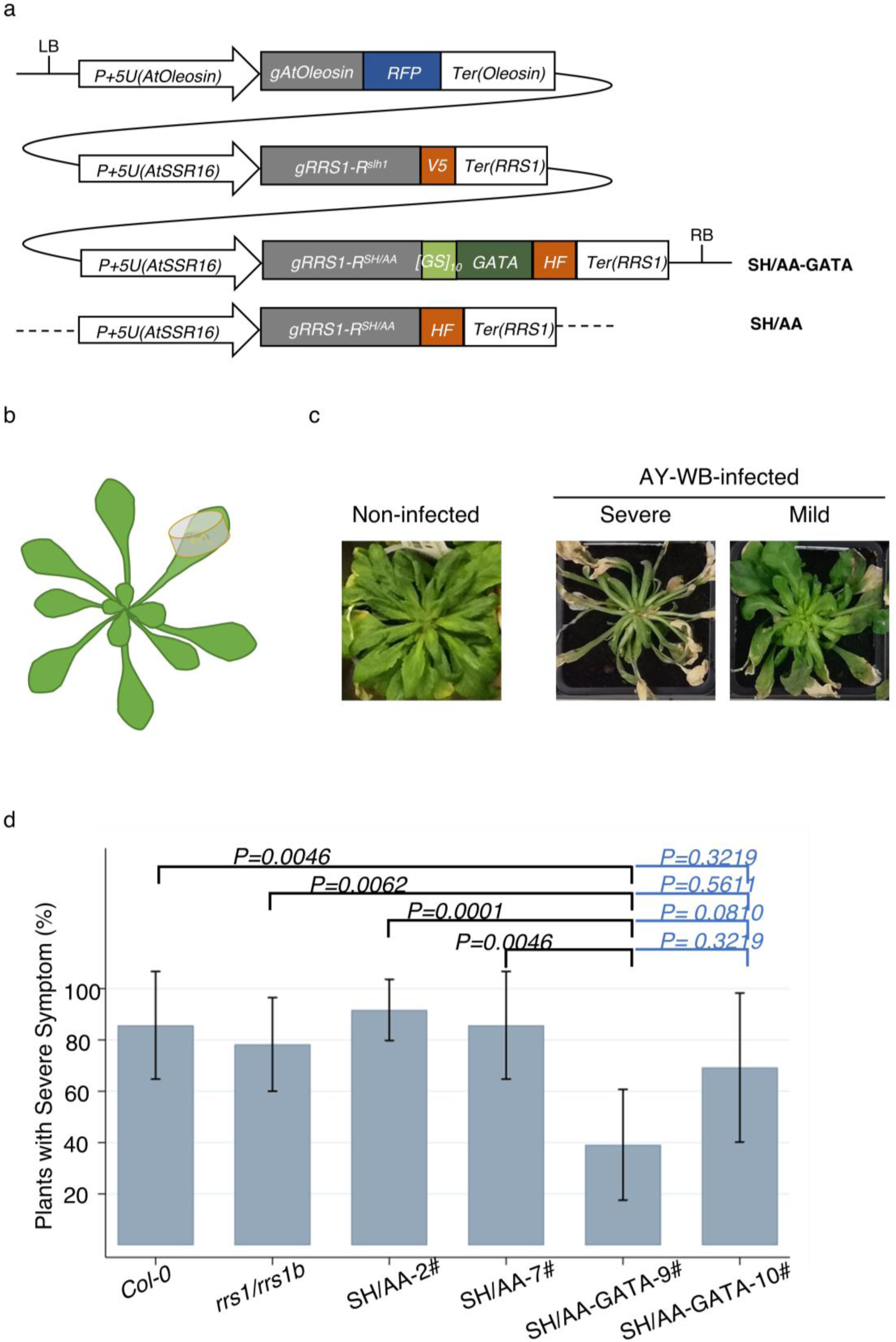
Transgenic Arabidopsis with RRS1-R^SH/AA^-[GS]_10_-GATA and RRS1-R^*slh1*^ has delayed disease symptoms after phytoplasma infection. **(a)** Schematic of constructs that were transformed into Arabidopsis. Arrow indicates the promoter, grey box indicates the gene, other coloured box indicates the linker region, epitope or GATA domain and terminators are in the last box. **(b)** Cartoon to show the phytoplasma infection. Leafhoppers carrying the phytoplasma were caged on one leaf for 3-4 days. **(c)** Disease development after phytoplasma infection. Infected Arabidopsis developed severe or mild symptoms 3-5 weeks post infection (wpi). **(d)** Statistical analysis of Arabidopsis 3-5 wpi with severe symptom. Infected Arabidopsis symptoms were characterised as severe or mild. Three independent infection experiments were performed, each line was tested for at least two times with the total number of plant n= 14, 23, 24, 14, 23 or 13. Student’s *t* test was used to analyse the difference level between groups. *P* value was indicated above the groups and error bar showed the 95% confidence interval). The analysis results are in Dataset_01.

Phytoplasmas are vectored by sap-feeding insects, including leafhopper, planthoppers and psyllids of the order Hemiptera^49^. We infected plants with AY-WB (Aster Yellows Witches’-Broom) phytoplasma carrying the SAP05 effector^35^ via transmission with aster leafhoppers (*Macrosteles quadrilineatus*) (Fig. 3b). The symptoms of AY-WB-infected Arabidopsis plants include small and deformed leaves, long petioles, pale green leaf colours, and narrow leaf blades^50,51^. After 3-5 weeks, some of the infected plants displayed severe symptoms whilst others displayed mild symptoms (Fig. 3c and S6). Seven to 10 weeks post infection (wpi), all the infected plants eventually died (Fig. S7).

We observed a delayed AY-WB phytoplasma disease symptom progression on infected SH/AA-GATA transgenic plants, compared to SH/AA, *Col-0/rrs1-3/rrs1b-1* or Col-0, 3-5 weeks after infection. The SH/AA-GATA-9# line shows a significant reduction in the number of plants with severe symptoms, while the reduction in line SH/AA-GATA-10# was not statistically significant (Fig. 3d). However, all infected plants including Col-0, *Col-0/rrs1-3/rrs1b-1*, SH/AA-2#, -7#, SH/AA-GATA-9#, and -10# died 7-10 weeks after phytoplasma infection (Fig. S7b), indicating that the transgenic Arabidopsis SH/AA-GATA are not fully resistant to AY-WB phytoplasma.

In this three-component system, the fusion protein RRS1-R^SH/AA^ -GATA suppressed the RPS4-dependent RRS1-R^*slh1*^ mediated autoactivity in the absence of effector. Upon effector recognition, the GATA domain fusion protein is degraded, restoring RRS1-R^*slh1*^ mediated autoactivity (Fig. 4). In the transient assay in *N. tabacum* and *N. benthamiana*, the HR occurs upon effector delivery (Fig. 2a). In transgenic Arabidopsis, this system delays the disease symptom progression upon phytoplasma infection (Fig. 3c and 3d), but the transgenic plants do not show full resistance.

**Figure 4.**
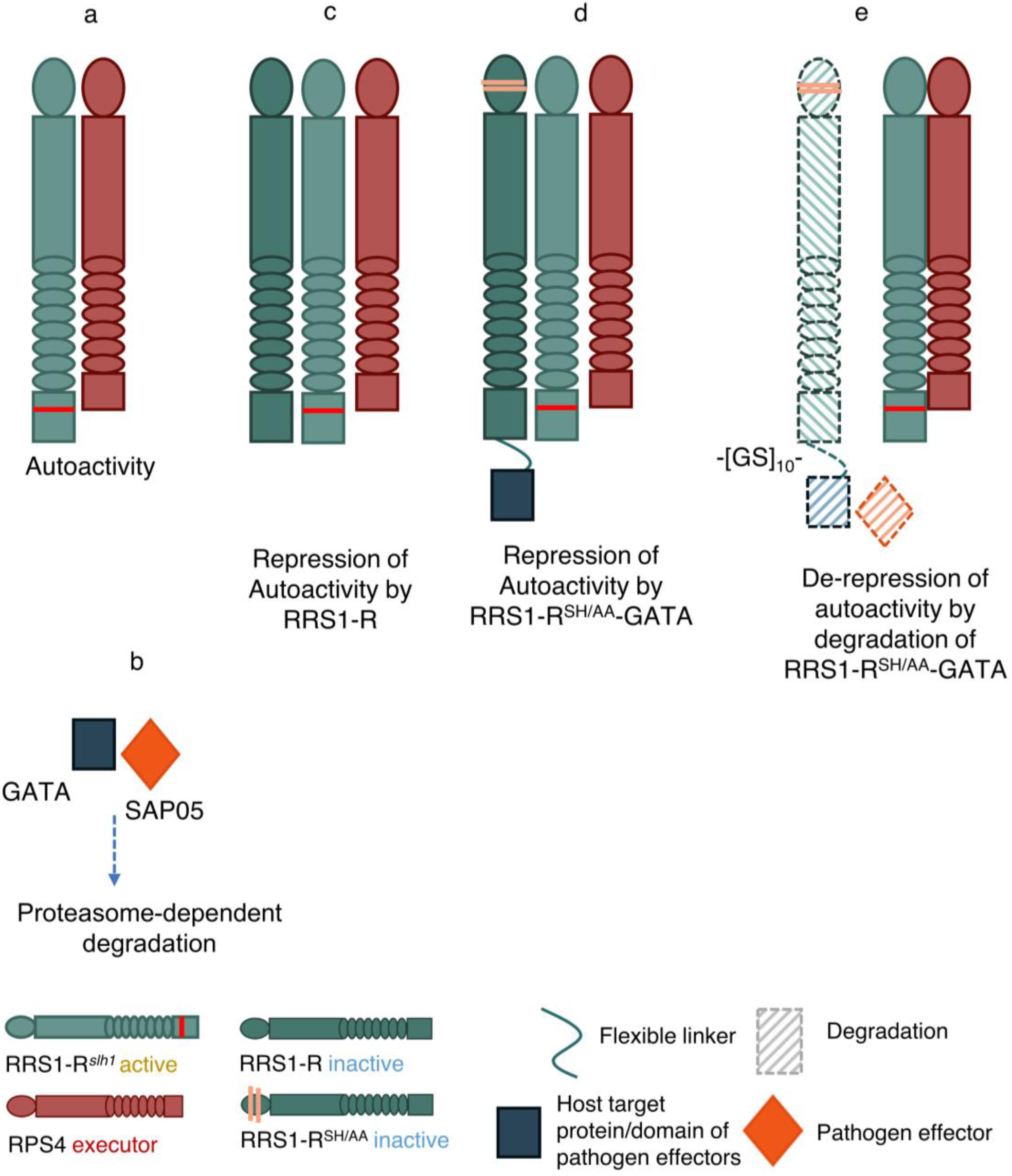
Schematic representation of engineering RRS1 with degron domain to recognize novel effector. **(a)** RRS1-R^*slh1*^ causes RPS4-dependent autoactivity. **(b)** Phytoplasma effector SAP05 targets GATA transcription factor and promotes proteasome-dependent degradation. **(c)** RRS1-R wildtype can repress RRS1-R^*slh1*^ autoactivity. **(d)** RRS1-R^*slh1*^ autoactivity can also be repressed by RRS1-R^SH/AA^-GATA. **(e)** The degradation of RRS1-R^SH/AA^-GATA by SAP05 derepresses RRS1-R^*slh1*^-induced-RPS4-dependent autoactivity.

## Discussion

Breeding crops to reduce losses to plant pathogens and pests involves judicious use of *Resistance* (*R*) genes, which usually encode NLR proteins that act as intracellular immune receptors which recognise effectors from pathogens and activate defense. Numerous NLR-encoding genes have been cloned, and there are widespread efforts to engineer known NLRs to expand their recognition capacity. We report here an approach to expanding the recognition capacity of the Arabidopsis paired NLRs RRS1 and RPS4. Transgenic tomato carrying RRS1-RPS4 show elevated resistance to *R. solanacearum* and *Pseudomonas syringae* pv. *tomato* DC3000 carrying the effector AvrRps4, and transgenic cucumber plants carrying RRS1/RPS4 are highly resistant to *Colletotrichum orbiculare*^52^. These data indicate that engineered RRS1/RPS4 can function in other species.

### Integrated domains as new source of resistance

The RRS1-R/RPS4 paired NLR recognises AvrRps4 and PopP2 via the integrated WRKY domain at the C terminus of RRS1-R. The discovery of non-canonical integrated domains in NLRs suggests a novel path for engineering NLR recognition capacity^6-8^. Rice NLR Pik-1 alleles which contain integrated HMA domains were designed to gain binding affinity and recognition capacity to the corresponding allelic effectors based on structural information or phylogenetic analysis. The modified Pik alleles activated immune responses in *N. benthamiana*^22-25^. Another HMA-containing rice NLR, RGA5 (which functions with RGA4), carrying an engineered HMA domain was able to perceive a new ligand in *N. benthamiana*^53^. In our system, RRS1-R^SH/AA^ -GATA recognizes an effector that has not been reported to be recognised by any NLR. Ideally, synthetic plant NLRs with any effector binding domain may result in new resistance genes. However, judging the best position to insert an ID is challenging^54^. Full length NLR structural studies will illuminate attempts to achieve such engineering. Here, for RRS1-R^SH/AA^-GATA, with the linker [GS]_10_ but not [NAAIR]_4_, can suppress RRS1-R^*slh1*^ autoactivity (Fig. S3), though this advance was achieved by trial-and-error rather than design.

### A novel approach for NLR engineering

Synthetic NLRs with modified HMA domains have been engineered with expanded recognition of allelic variants of effectors^22-25^. However, further enhancement of the effector spectrum for NLR engineering would be desirable. With more knowledge of effectors and host targets, new strategies are emerging for synthetic NLRs that can respond to divergent effectors. In the Arabidopsis PBS1/RPS5 decoy system, PBS1 contains a protease cleavage site that serves as decoy and NLR RPS5 is activated by proteolytic cleavage of PBS1. This system mediates detection of the effector protein AvrPphB from *Pseudomonas syringae*^19^. By swapping the proteolytic target site of PBS1 to protease recognition sites of AvrRpt2 from *P. syringae* or NIa protease from TEV or TuMV, the engineered PBS1 with RPS5 confers resistance to *P. syringae* carring AvrRpt2, TEV or TuMV^20^. In our system, we derepress RRS1-R^*slh1*^ autoimmunity by attaching an effector-dependent degron to inactive, repressing RRS1-R^*slh1*^. We generated the chimeric repressor RRS1-R^SH/AA^-GATA to be degraded in response to a phytoplasma effector SAP05, thus broadening the NLR recognition specificity. Incorporating SH/AA mutation in the signalling TIR domain was required to prevent the autoactivity of the fusion protein. With the expanding knowledge of NLRs and effectors, additional alternative strategies to design synthetic NLRs will enable further engineering to expand NLR recognition specificity.

### HR is not always associated with resistance

HR is a form of localized cell death that occurs at the pathogen infection sites, and is often correlated recognition of a pathogen effector by a plant R protein^55^. Recent data show how surface receptor- and intracellular receptor-mediated immunity mechanisms mutually potentiate, culminating in an HR^56,57^. In plant-virus pathosystems, resistance can occur in the absence of cell death. Potato *Rx* mediates extreme resistance to various Potato Virus X (PVX) strains in *Nicotiana* spp and potato, without visible cell death^58^. Transgenic *N. benthamiana* plants carrying the tomato *Tm-2*^*2*^ gene show extreme resistance to Tobacco Mosaic Virus (TMV) in the absence of cell death^59^. In the plant-oomycete pathosystem, potato *Rpi-chc1*.*2* confers HR on coexpression with multiple *Phytophthora infestans* effector PexRD31 paralogs, but doesn’t always confer late blight resistance^60^. The lack of full correlation of cell death and resistance is also observed in Arabidopsis and bacterial pathogens. The Arabidopsis *nrg1* mutant lacks HR induced by RRS1/RPS4 and RRS1B/RPS4B but still maintains the resistance to *Pseudomonas syringae* pv. *tomato* strain DC3000 carrying effector AvrRps4 or AvrRpt2 (recognised by CC-NLR RPS2)^61^. PAD4 is dispensable for RRS1/RPS4 mediated cell death but required for the resistance to DC3000 with AvrRps4^62^. RGA5 with a modified HMA domain can confer HR to a new ligand, AVR-PikD in *N. benthamiana* but does not confer resistance in the homologous rice system ^53^.

Despite effector-dependent HR in transient assays in *Nicotiana* sp., transgenic Arabidopsis carrying RPS4, RRS1-R^*slh1*^ and RRS1-R^SH/AA^-GATA were not resistant to Phytoplasma infection, though they did show delayed symptoms. This further exemplifies how HR does not always correlate to resistance. Phytoplasmas locate to the phloem and it remains an open question whether NLRs induce HR in all cell types. Also, whereas SAP05 is likely to unload from the phloem and migrate to other cells, it is not yet clear which cells the effector migrates to and whether these cells have the signaling pathway that translates NLR activation into an HR. Finally, SAP05 degrades plant transcription factors and changes developmental process^35^ and may interfere with defense induction upstream of NLR signaling. Conceivably, there is insufficient SAP05 delivery from phytoplasma infection to enable enough RRS1-R^SH/AA^-GATA degradation to fully derepress immunity. Caution is required in selecting the appropriate effectors for NLR engineering. It is likely that a balance needs to be struck in our system between expressing sufficient RRS1-R^SH/AA^-GATA to avoid any constitutive defense activation by RRS1-R^*slh1*^, while avoiding expressing so much RRS1-R^*slh1*^ that a pathogen effector fails to remove enough to derepress RRS1-R^*slh1*^.

TNL-mediated cell death and resistance requires a non-NLR protein, Enhanced Disease Susceptibility1 (EDS1) in Arabidopsis. EDS1 forms EDS1-SAG101 or EDS1-PAD4 complexes to determine the cell death and pathogen growth restriction respectively^63,64^, and EDS1-SAG101 engages with the RPW8-NLR NRG1 upon effector recognition^65^. Since the engineered NLR is RPS4-dependent, it is likely to signal via these modules. NLR protein levels also determine the strength of resistance^59^; R genes are usually semi-dominant, and transgenic lines with sufficient NLR expression are likely required for robust resistance. Last but not least, engineering the resistance to a phloem-limited insect-transmitted plant pathogen might require more knowledge of defense in phloem.

In summary, this study provides a new strategy utilizing the concept of integrated domain and the recessiveness of RRS1 to expand NLR-dependent effector recognition capacity.

## Acknowledgements

S.W., Z.D. were supported on ERC grant “Immunitybypairdesign” Project ID 669926 to JDGJ. W.H. was supported by the Human Frontier Science Program RGP0024/2105. Additional support was received from Biotechnology and Biological Sciences Research Council grants BB/K002848/1, BB/J0045531/1 and BB/P012574/1 and the John Innes Foundation.

We thank the tissue culture team in The Sainsbury Laboratory for generating the transgenic Arabidopsis plants and the JIC Entomology Facility for rearing of leafhopper and phytoplasma stocks. We thank Dr Agnieszka Alexander for her diligent work and support as a lab manager from Jones Lab. We thank Vicenç Esteve-Guasch from University of East Anglia for the discussion of statistical analysis.

## Contribution

J.D.G.J. and S.A.H. conceptualized the study and obtained funding. S.W. and J.D.G.J. designed the experiment. S.W., W.H. and Z.D. performed the experiments and data analysis. S.W. and J.D.G.J. wrote the manuscript. All the authors edited the manuscript.

## Supplementary Materials

Materials and Methods

Figs. S1 to S7

Table S1 to S4

## Supplementary Figure

**Fig. S1.**
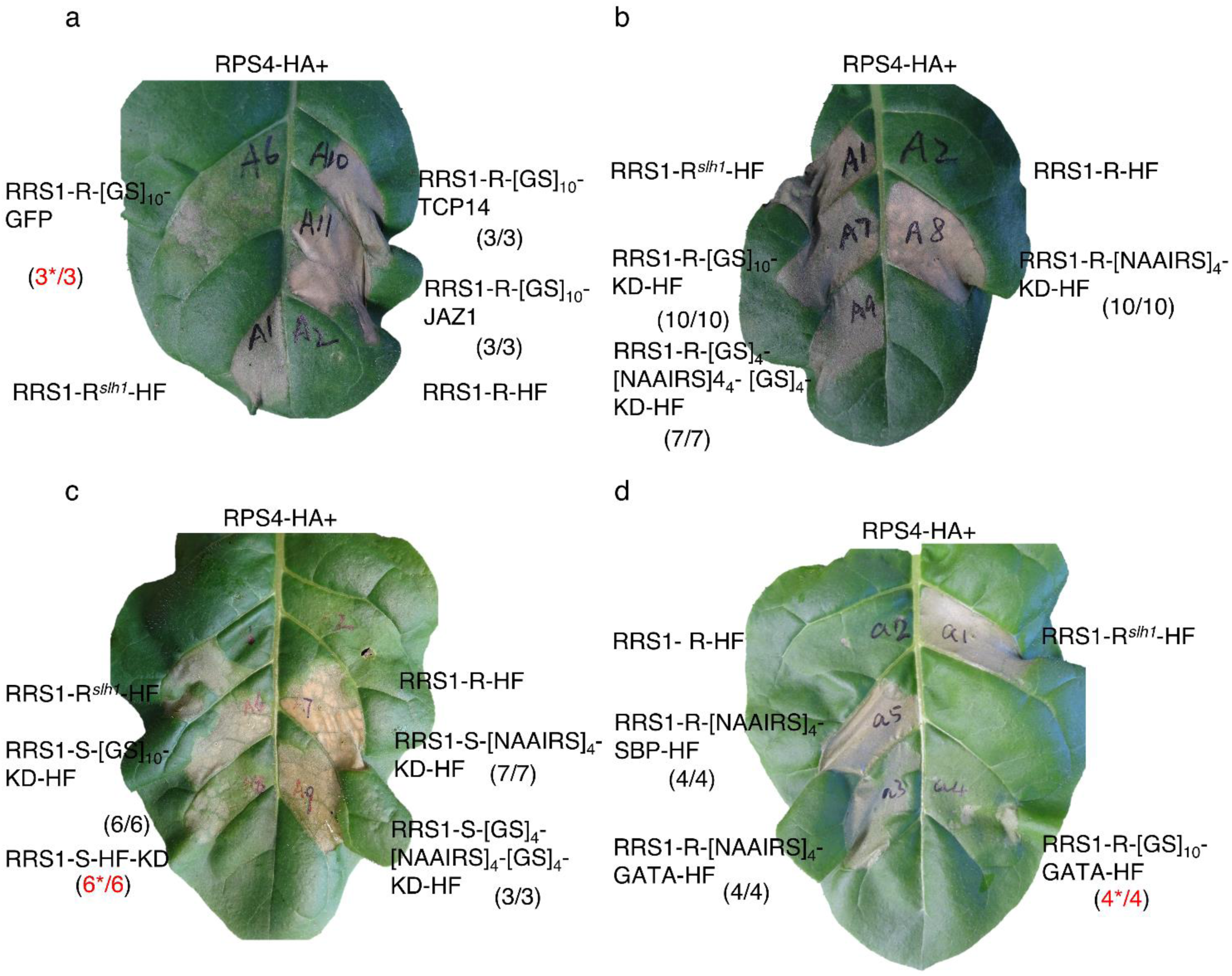
HR assays to test the RPS4-dependent autoactivity of RRS1-X constructs. *N. tabacum* leaves were co-infiltrated with *A. tumefaciens* strains (each at OD_600_ = 0.5) carrying RPS4 and the indicating RRS1-X construct. **(a)** RPS4 and RRS1-R-[GS]_10_-GFP, RRS1-R-[GS]_10_-TCP14, RRS1-R-[GS]_10_-JAZ1, showed RPS4-dependent autoactivity. **(b)** RRS1-R and KD fusion protein with three different linkers ([GS]_10_, [NAAIRS]_4_ and [GS]_10_-[NAAIRS]_4_-[GS]_10_) all displayed RPS4-dependent autoactivity. **(c)** RRS1-S and KD fusion protein with 4 different linkers ([GS]_10_, [NAAIRS]_4_, [GS]_10_-[NAAIRS]_4_-[GS]_10_ and HF) all displayed RPS4-dependent autoactivity. **(d)** RRS1-R-[NAAIRS]_4_-SBP-HF and RRS1-R-[NAAIRS]_4_-GATA-HF displayed strong RPS4-dependent autoactivity whilst RRS1-R-[GS]_10_-GATA-HF displayed weak RPS4-dependent autoactivity. HR was visually assessed and photographed at 3 days post-infiltration (3 dpi). When co-infiltrated with RPS4, RRS1-R^*slh1*^-HF (HR) and RRS1-R-HF (no HR) were used as control. The numbers in parentheses are the number of leaves displaying HR out of the total number of leaves infiltrated. Numbers in red with ‘^*^’ represents weak HR.

**Fig. S2.**
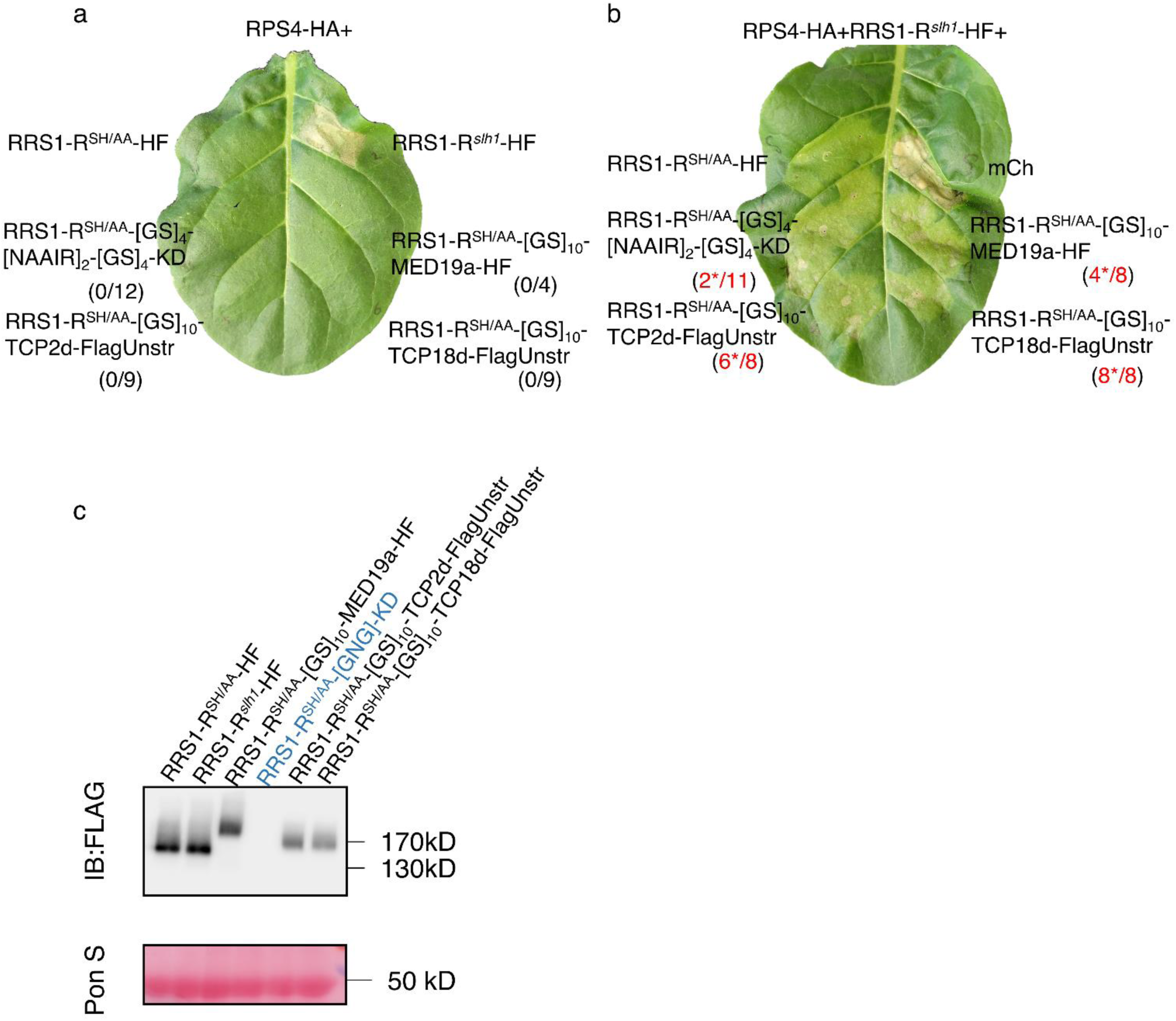
RRS1-R^SH/AA^-X constructs do not show RPS4-dependent autoactivity but cannot fully suppress RRS1-R^*slh1*^ mediated-autoactivity. **(a)** *N. tabacum* leaves were co-infiltrated with *A. tumefaciens* strains (each at OD_600_ = 0.5) carrying RPS4 and RRS1-R^SH/AA^-[GS]_4_-[NAAIR]_2_-[GS]_4_-KD-HF, RRS1-R^SH/AA^-[GS]_10_-MED19a-HF, RRS1-R^SH/AA^-[GS]_10_-TCP2d-HF or RRS1-R^SH/AA^-[GS]_10_-TCP18d-HF did not show HR. **(b)** All the RRS1-R^SH/AA^-X constructs tested in (A) cannot fully repress RRS1-R^*slh1*^-mediated autoactivity. The numbers in parentheses are the number of leaves displaying HR out of the total number of leaves infiltrated. Numbers in red with ‘^*^’ represents partial cell death. **(c)** Western blot showed the protein expression level. *N. benthamiana* leaves were infiltrated with *A. tumefaciens* strains (OD_600_= 0.5) carrying RRS1-R^SH/AA^-[GS]_4_-[NAAIR]_2_-[GS]_4_-KD-HF, RRS1-R^SH/AA^-[GS]_10_-MED19a-HF, RRS1-R^SH/AA^-[GS]_10_-TCP2d-FlagUnstr or RRS1-R^SH/AA^-[GS]_10_-TCP18d-FlagUnstr. Samples were harvested 48 hours post infiltration (hpi) and total protein were extracted and subjected to SDS-PAGE, then immunoblotted for FLAG. The construct highlighted in blue has no FLAG epitope. Ponceau S staining (Pon S) was used as loading control.

**Fig. S3.**
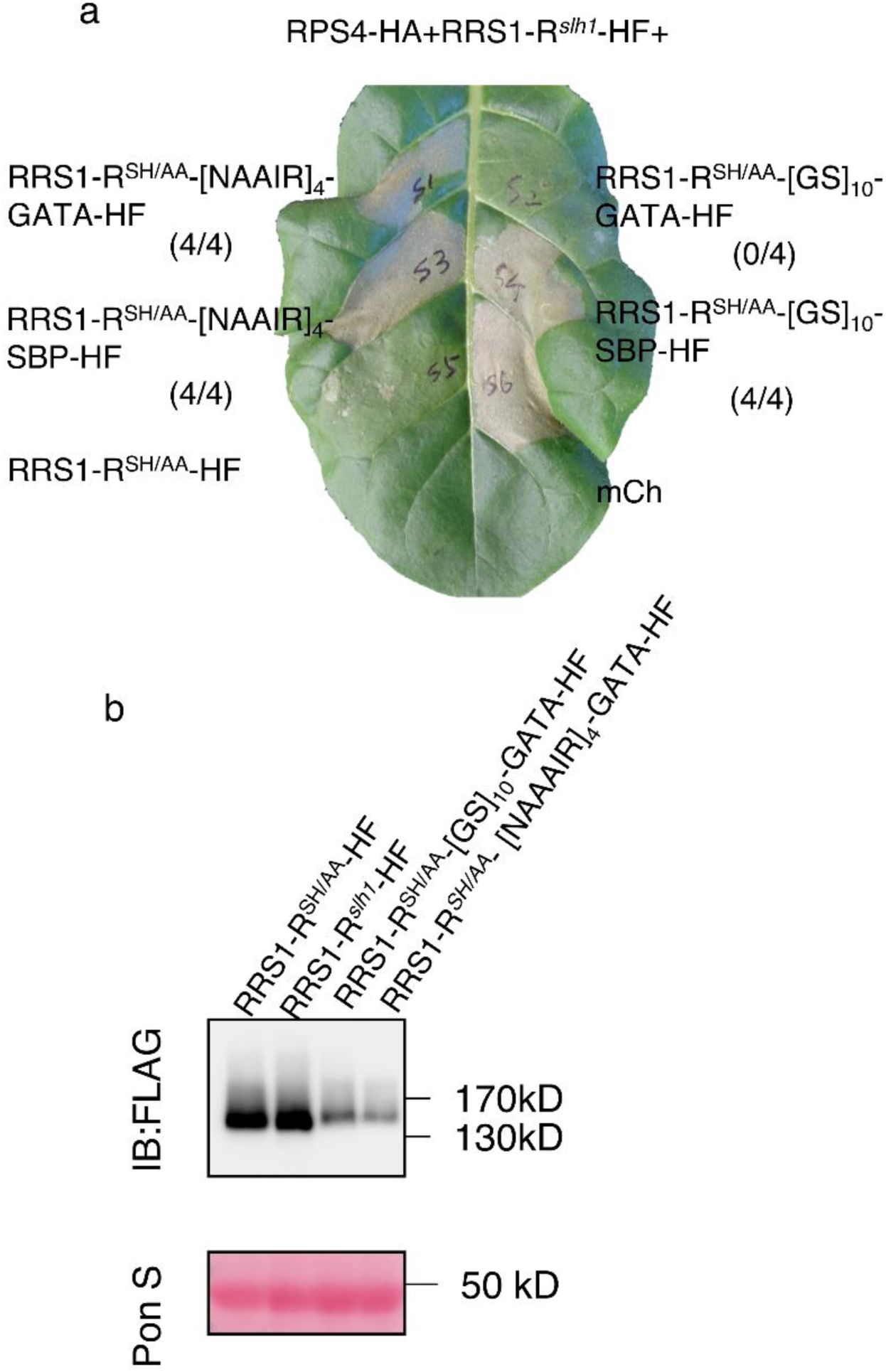
The linker between RRS1-R^SH/AA^ and fusion domain is important to suppress RRS1-R^*slh1*^-induced-autoactivity. **(a)** *N. tabacum* leaves were co-infiltrated with *A. tumefaciens* strains (each at OD_600_ = 0.5) carrying RPS4, RRS1-R^*slh1*^and RRS1-R^SH/AA^-[NAAIR]_4_-GATA-HF, RRS1-R^SH/AA^- [GS]_10_-GATA-HF, RRS1-R^SH/AA^-[NAAIR]_4_-SBP-HF or RRS1-R^SH/AA^-[GS]_10_-SBP-HF. Only RRS1-R^SH/AA^-[GS]_10_-GATA-HF can fully suppress RRS1-R^*slh1*^ mediated-autoactivity. **(b)** Western blot is to show the protein expression of RRS1-R^SH/AA^-[NAAIR]_4_-GATA-HF and RRS1-R^SH/AA^-[GS]_10_-GATA-HF. The numbers in parentheses are the number of leaves displaying HR out of the total number of leaves infiltrated.

**Fig. S4.**
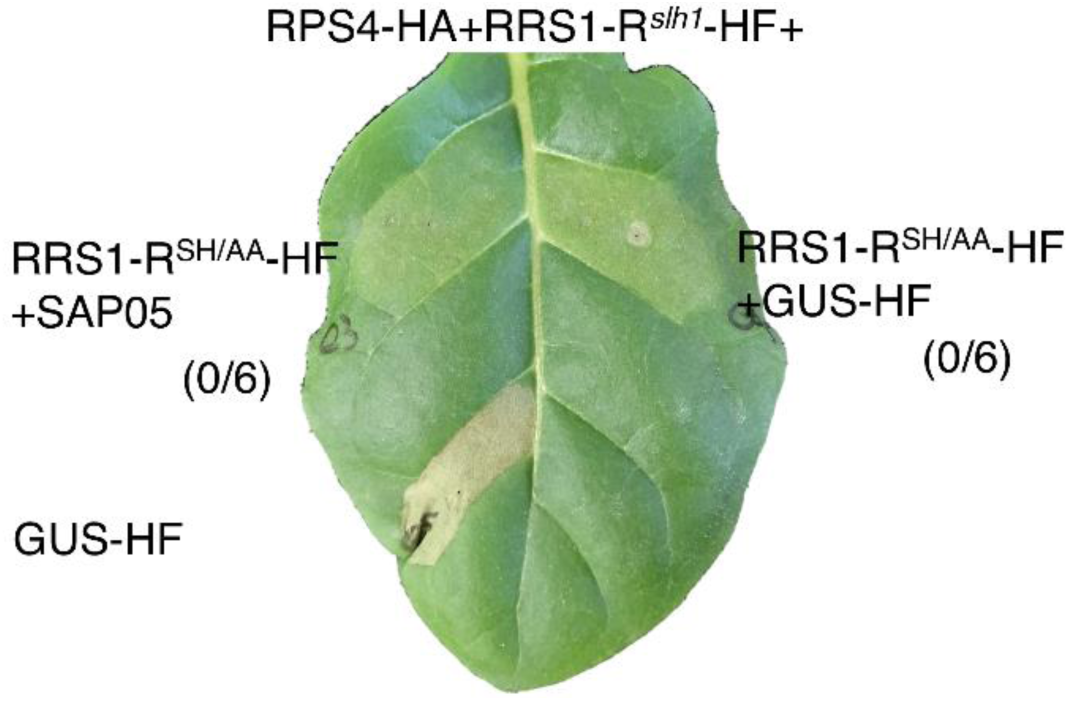
RRS1-R^SH/AA^ does not respond to SAP05. *N. tabacum* leaves were co-infiltrated with *A. tumefaciens* strains (each at OD_600_ = 0.5, strain with SAP05 OD_600_ = 1) carrying RPS4, RRS1-R^*slh1*^, RRS1-R^SH/AA^-HF and SAP05 or GUS-HF. RRS1-R^SH/AA^-HF did not show cell death when co-expressed with SAP05. The numbers in parentheses are the number of leaves displaying HR out of the total number of leaves infiltrated.

**Fig. S5.**
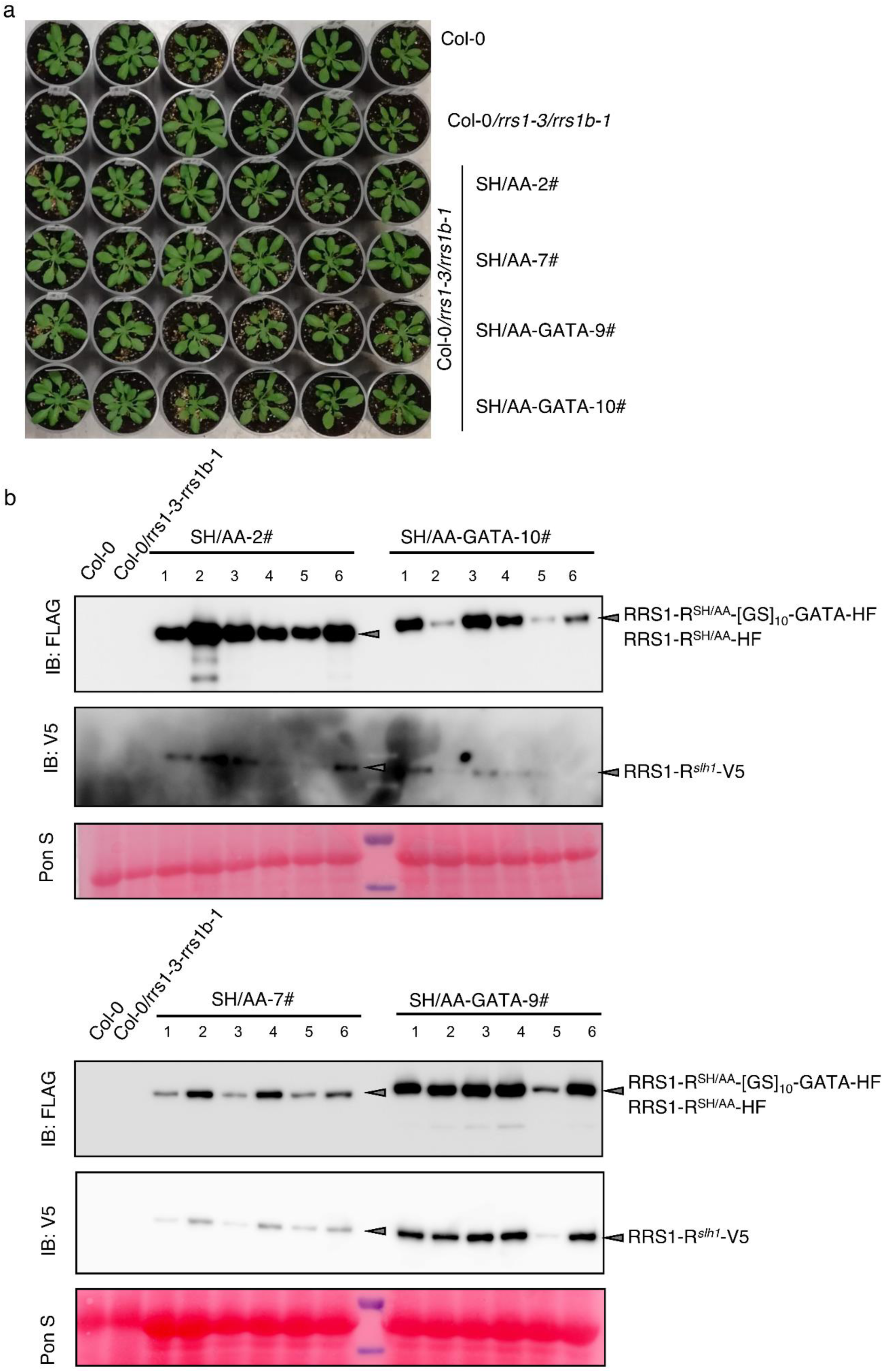
Phenotype and protein expression level of transgenic Arabidopsis. **(a)** Phenotype of the indicated 3-4 weeks-old *Arabidopsis* plants. Transgenic Arabidopsis lines (T2 plants) transformed with RRS1-R^*slh1*^ -V5 and RRS1-R^SH/AA^-HF or RRS1-R^SH/AA^-[GS]_10_-GATA-HF were labled as SH/AA or SH/AA-GATA respectively. **(b)** Western blot results to show the protein expression level of each transgenic line. Arabidopsis leaves were harvested from 3-4 weeks-old plants individually. Total protein was extracted and subjected to SDS-PAGE and immunoblotted for FLAG or V5. Pon S was used as loading control.

**Fig. S6.**
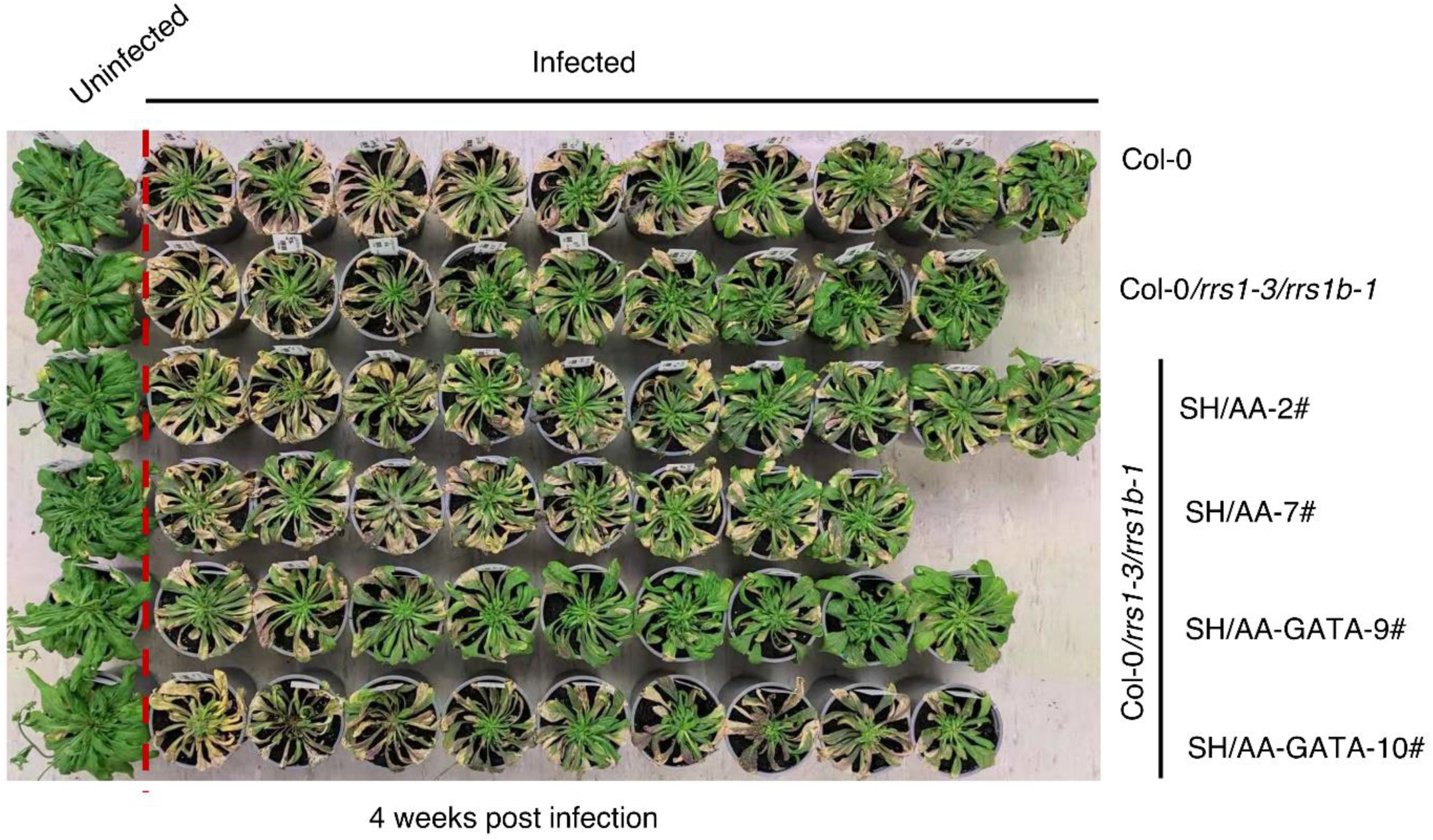
Disease symptom of Arabidopsis 4 weeks post phytoplasma infection.

**Fig. S7.**
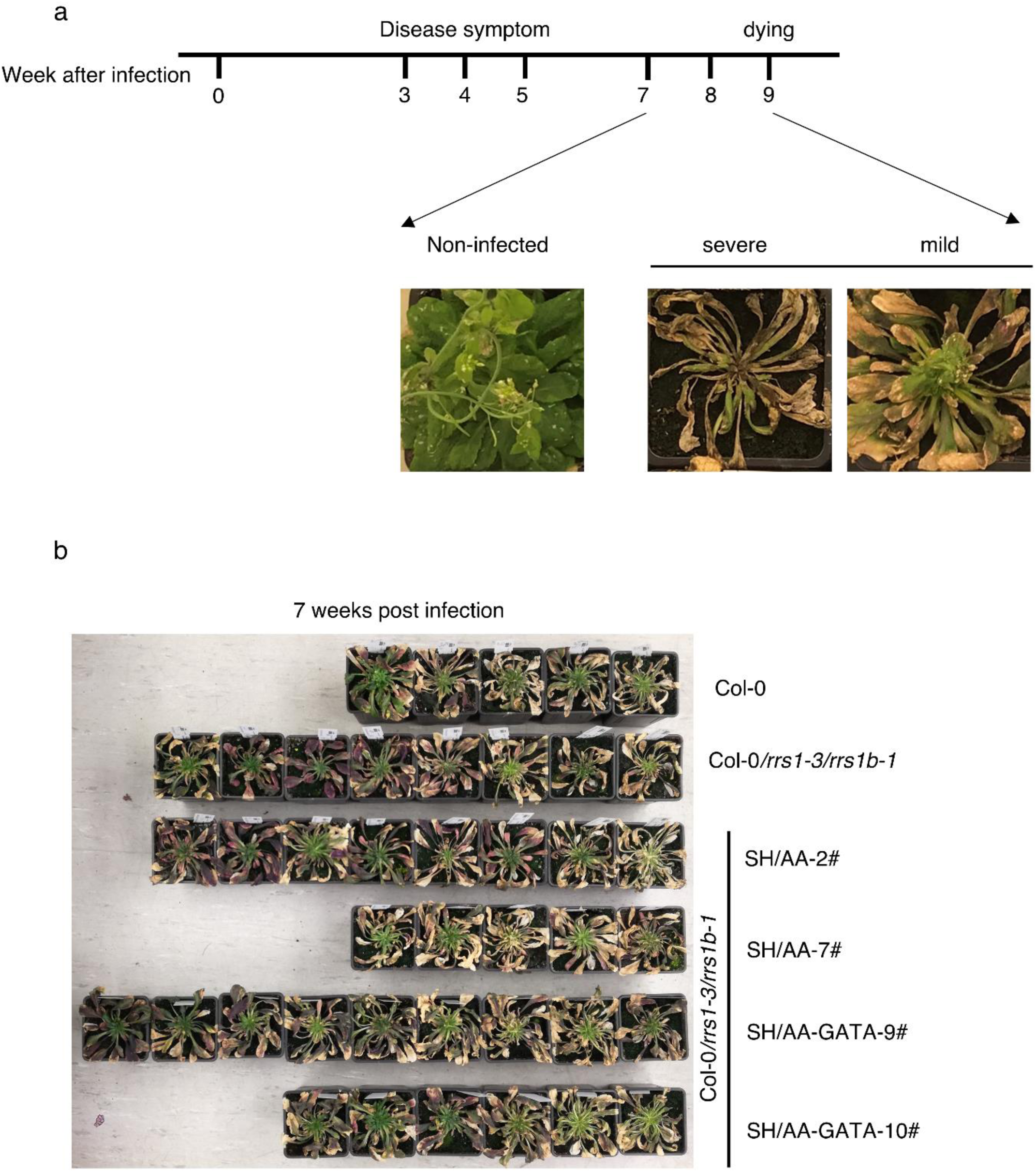
Disease symptom of Arabidopsis in the late stage after phytoplasma infection. **(a)** Timeline of infection assay and typical symptom of Arabidopsis plant 7-9 weeks post phytoplasma infection. **(b)** Arabidopsis were dying 7 weeks post phytoplasma infection.

## References

1 Jones, J. D. & Dangl, J. L. The plant immune system. Nature 444, 323–329, doi:10.1038/nature05286 (2006).

2 Jones, J. D., Vance, R. E. & Dangl, J. L. Intracellular innate immune surveillance devices in plants and animals. Science 354, doi:10.1126/science.aaf6395 (2016).

3 Whitham, S. et al. The product of the tobacco mosaic virus resistance gene N: similarity to toll and the interleukin-1 receptor. Cell 78, 1101–1115, doi:10.1016/0092-8674(94)90283-6 (1994).

4 Bent, A. F. et al. RPS2 of Arabidopsis thaliana: a leucine-rich repeat class of plant disease resistance genes. Science 265, 1856–1860, doi:10.1126/science.8091210 (1994).

5 Kourelis, J. & van der Hoorn, R. A. L. Defended to the Nines: 25 Years of Resistance Gene Cloning Identifies Nine Mechanisms for R Protein Function. Plant Cell 30, 285–299, doi:10.1105/tpc.17.00579 (2018).

6 Kroj, T., Chanclud, E., Michel-Romiti, C., Grand, X. & Morel, J. B. Integration of decoy domains derived from protein targets of pathogen effectors into plant immune receptors is widespread. New Phytol 210, 618–626, doi:10.1111/nph.13869 (2016).

7 Sarris, P. F., Cevik, V., Dagdas, G., Jones, J. D. & Krasileva, K. V. Comparative analysis of plant immune receptor architectures uncovers host proteins likely targeted by pathogens. BMC Biol 14, 8, doi:10.1186/s12915-016-0228-7 (2016).

8 Bailey, P. C. et al. Dominant integration locus drives continuous diversification of plant immune receptors with exogenous domain fusions. Genome Biol 19, 23, doi:10.1186/s13059-018-1392-6 (2018).

9 Van de Weyer, A. L. et al. A Species-Wide Inventory of NLR Genes and Alleles in Arabidopsis thaliana. Cell 178, 1260–1272 e1214, doi:10.1016/j.cell.2019.07.038 (2019).

10 Zhang, L. et al. Cryo-EM structure of the activated NAIP2-NLRC4 inflammasome reveals nucleated polymerization. Science 350, 404–409, doi:10.1126/science.aac5789 (2015).

11 Wang, J. et al. Reconstitution and structure of a plant NLR resistosome conferring immunity. Science 364, doi:10.1126/science.aav5870 (2019).

12 Wang, J. et al. Ligand-triggered allosteric ADP release primes a plant NLR complex. Science 364, doi:10.1126/science.aav5868 (2019).

13 Ma, S. et al. Direct pathogen-induced assembly of an NLR immune receptor complex to form a holoenzyme. Science 370, doi:10.1126/science.abe3069 (2020).

14 Duxbury, Z. et al. Induced proximity of a TIR signaling domain on a plant-mammalian NLR chimera activates defense in plants. Proc Natl Acad Sci U S A 117, 18832–18839, doi:10.1073/pnas.2001185117 (2020).

15 Farnham, G. & Baulcombe, D. C. Artificial evolution extends the spectrum of viruses that are targeted by a disease-resistance gene from potato. Proc Natl Acad Sci U S A 103, 18828–18833, doi:10.1073/pnas.0605777103 (2006).

16 Segretin, M. E. et al. Single amino acid mutations in the potato immune receptor R3a expand response to Phytophthora effectors. Mol Plant Microbe Interact 27, 624–637, doi:10.1094/MPMI-02-14-0040-R (2014).

17 Giannakopoulou, A. et al. Tomato I2 Immune Receptor Can Be Engineered to Confer Partial Resistance to the Oomycete Phytophthora infestans in Addition to the Fungus Fusarium oxysporum. Mol Plant Microbe Interact 28, 1316–1329, doi:10.1094/MPMI-07-15-0147-R (2015).

18 Bernoux, M. et al. Comparative Analysis of the Flax Immune Receptors L6 and L7 Suggests an Equilibrium-Based Switch Activation Model. Plant Cell 28, 146–159, doi:10.1105/tpc.15.00303 (2016).

19 Simonich, M. T. & Innes, R. W. A disease resistance gene in Arabidopsis with specificity for the avrPph3 gene of Pseudomonas syringae pv. phaseolicola. Mol Plant Microbe Interact 8, 637–640, doi:10.1094/mpmi-8-0637 (1995).

20 Kim, S. H., Qi, D., Ashfield, T., Helm, M. & Innes, R. W. Using decoys to expand the recognition specificity of a plant disease resistance protein. Science 351, 684–687, doi:10.1126/science.aad3436 (2016).

21 Maqbool, A. et al. Structural basis of pathogen recognition by an integrated HMA domain in a plant NLR immune receptor. Elife 4, doi:10.7554/eLife.08709 (2015).

22 De la Concepcion, J.C. et al. Protein engineering expands the effector recognition profile of a rice NLR immune receptor. Elife 8, doi:10.7554/eLife.47713 (2019).

23 De la Concepcion, J.C. et al. Polymorphic residues in rice NLRs expand binding and response to effectors of the blast pathogen. Nat Plants 4, 576–585, doi:10.1038/s41477-018-0194-x (2018).

24 De la Concepcion, J.C. et al. The allelic rice immune receptor Pikh confers extended resistance to strains of the blast fungus through a single polymorphism in the effector binding interface. PLoS Pathog 17, e1009368, doi:10.1371/journal.ppat.1009368 (2021).

25 Białas, A. et al. Two NLR immune receptors acquired high-affinity binding to a fungal effector through convergent evolution of their integrated domain. bioRxiv, 2021.2001.2026.428286, doi:10.1101/2021.01.26.428286 (2021).

26 Narusaka, M. et al. RRS1 and RPS4 provide a dual Resistance-gene system against fungal and bacterial pathogens. Plant J 60, 218–226, doi:10.1111/j.1365-313X.2009.03949.x (2009).

27 Gassmann, W., Hinsch, M. E. & Staskawicz, B. J. The Arabidopsis RPS4 bacterial-resistance gene is a member of the TIR-NBS-LRR family of disease-resistance genes. Plant J 20, 265–277, doi:10.1046/j.1365-313x.1999.t01-1-00600.x (1999).

28 Deslandes, L. et al. Physical interaction between RRS1-R, a protein conferring resistance to bacterial wilt, and PopP2, a type III effector targeted to the plant nucleus. Proc Natl Acad Sci U S A 100, 8024–8029, doi:10.1073/pnas.1230660100 (2003).

29 Le Roux, C. et al. A receptor pair with an integrated decoy converts pathogen disabling of transcription factors to immunity. Cell 161, 1074–1088, doi:10.1016/j.cell.2015.04.025 (2015).

30 Sarris, P. F. et al. A Plant Immune Receptor Detects Pathogen Effectors that Target WRKY Transcription Factors. Cell 161, 1089–1100, doi:10.1016/j.cell.2015.04.024 (2015).

31 Ma, Y. et al. Distinct modes of derepression of an Arabidopsis immune receptor complex by two different bacterial effectors. Proc Natl Acad Sci U S A 115, 10218–10227, doi:10.1073/pnas.1811858115 (2018).

32 Guo, H. et al. Phosphorylation-Regulated Activation of the Arabidopsis RRS1-R/RPS4 Immune Receptor Complex Reveals Two Distinct Effector Recognition Mechanisms. Cell Host Microbe 27, 769–781 e766, doi:10.1016/j.chom.2020.03.008 (2020).

33 Noutoshi, Y. et al. A single amino acid insertion in the WRKY domain of the Arabidopsis TIR-NBS-LRR-WRKY-type disease resistance protein SLH1 (sensitive to low humidity 1) causes activation of defense responses and hypersensitive cell death. Plant J 43, 873–888, doi:10.1111/j.1365-313X.2005.02500.x (2005).

34 Sohn, K. H. et al. The nuclear immune receptor RPS4 is required for RRS1SLH1-dependent constitutive defense activation in Arabidopsis thaliana. PLoS Genet 10, e1004655, doi:10.1371/journal.pgen.1004655 (2014).

35 Huang, W. et al. Parasite co-opts a ubiquitin receptor to induce a plethora of developmental changes. bioRxiv, 2021.2002.2015.430920, doi:10.1101/2021.02.15.430920 (2021).

36 Pecher, P. et al. Phytoplasma SAP11 effector destabilization of TCP transcription factors differentially impact development and defence of Arabidopsis versus maize. PLoS Pathog 15, e1008035, doi:10.1371/journal.ppat.1008035 (2019).

37 Sugio, A., Kingdom, H. N., MacLean, A. M., Grieve, V. M. & Hogenhout, S. A. Phytoplasma protein effector SAP11 enhances insect vector reproduction by manipulating plant development and defense hormone biosynthesis. Proc Natl Acad Sci U S A 108, E1254–1263, doi:10.1073/pnas.1105664108 (2011).

38 Sugio, A., MacLean, A. M. & Hogenhout, S. A. The small phytoplasma virulence effector SAP11 contains distinct domains required for nuclear targeting and CIN-TCP binding and destabilization. New Phytol 202, 838–848, doi:10.1111/nph.12721 (2014).

39 MacLean, A. M. et al. Phytoplasma effector SAP54 hijacks plant reproduction by degrading MADS-box proteins and promotes insect colonization in a RAD23-dependent manner. PLoS Biol 12, e1001835, doi:10.1371/journal.pbio.1001835 (2014).

40 Yang, L. et al. Pseudomonas syringae Type III Effector HopBB1 Promotes Host Transcriptional Repressor Degradation to Regulate Phytohormone Responses and Virulence. Cell Host Microbe 21, 156–168, doi:10.1016/j.chom.2017.01.003 (2017).

41 Gimenez-Ibanez, S. et al. The bacterial effector HopX1 targets JAZ transcriptional repressors to activate jasmonate signaling and promote infection in Arabidopsis. PLoS Biol 12, e1001792, doi:10.1371/journal.pbio.1001792 (2014).

42 Mosher, R. A., Durrant, W. E., Wang, D., Song, J. & Dong, X. A comprehensive structure-function analysis of Arabidopsis SNI1 defines essential regions and transcriptional repressor activity. Plant Cell 18, 1750–1765, doi:10.1105/tpc.105.039677 (2006).

43 Wilson, I. A. et al. Identical short peptide sequences in unrelated proteins can have different conformations: a testing ground for theories of immune recognition. Proc Natl Acad Sci U S A 82, 5255–5259, doi:10.1073/pnas.82.16.5255 (1985).

44 Williams, S. J. et al. Structural basis for assembly and function of a heterodimeric plant immune receptor. Science 344, 299–303, doi:10.1126/science.1247357 (2014).

45 Caillaud, M. C. et al. A downy mildew effector attenuates salicylic acid-triggered immunity in Arabidopsis by interacting with the host mediator complex. PLoS Biol 11, e1001732, doi:10.1371/journal.pbio.1001732 (2013).

46 Fishbain, S., Prakash, S., Herrig, A., Elsasser, S. & Matouschek, A. Rad23 escapes degradation because it lacks a proteasome initiation region. Nat Commun 2, 192, doi:10.1038/ncomms1194 (2011).

47 Engler, C. et al. A golden gate modular cloning toolbox for plants. ACS Synth Biol 3, 839–843, doi:10.1021/sb4001504 (2014).

48 Shimada, T. L., Shimada, T. & Hara-Nishimura, I. A rapid and non-destructive screenable marker, FAST, for identifying transformed seeds of Arabidopsis thaliana. Plant J 61, 519–528, doi:10.1111/j.1365-313X.2009.04060.x (2010).

49 Jiang, Y., Zhang, C. X., Chen, R. & He, S. Y. Challenging battles of plants with phloem-feeding insects and prokaryotic pathogens. Proc Natl Acad Sci U S A 116, 23390–23397, doi:10.1073/pnas.1915396116 (2019).

50 MacLean, A. M. et al. Phytoplasma effector SAP54 induces indeterminate leaf-like flower development in Arabidopsis plants. Plant Physiol 157, 831–841, doi:10.1104/pp.111.181586 (2011).

51 Hogenhout, S. A. et al. Phytoplasmas: bacteria that manipulate plants and insects. Mol Plant Pathol 9, 403–423, doi:10.1111/j.1364-3703.2008.00472.x (2008).

52 Narusaka, M. et al. Interfamily transfer of dual NB-LRR genes confers resistance to multiple pathogens. PLoS One 8, e55954, doi:10.1371/journal.pone.0055954 (2013).

53 Cesari, S. et al. Design of a new effector recognition specificity in a plant NLR immune receptor by molecular engineering of its integrated decoy domain. bioRxiv (2021).

54 Ellis, J. G. Integrated decoys and effector traps: how to catch a plant pathogen. BMC Biol 14, 13, doi:10.1186/s12915-016-0235-8 (2016).

55 Flor, H. H. Current Status of the Gene-For-Gene Concept. Annual Review of Phytopathology 9, 275–296, doi:10.1146/annurev.py.09.090171.001423 (1971).

56 Ngou, B. P. M., Ahn, H. K., Ding, P. & Jones, J. D. G. Mutual potentiation of plant immunity by cell-surface and intracellular receptors. Nature 592, 110–115, doi:10.1038/s41586-021-03315-7 (2021).

57 Yuan, M. et al. Pattern-recognition receptors are required for NLR-mediated plant immunity. Nature 592, 105–109, doi:10.1038/s41586-021-03316-6 (2021).

58 Bendahmane, A., Kanyuka, K. & Baulcombe, D. C. The Rx gene from potato controls separate virus resistance and cell death responses. Plant Cell 11, 781–792, doi:10.1105/tpc.11.5.781 (1999).

59 Zhang, H., Zhao, J., Liu, S., Zhang, D. P. & Liu, Y. Tm-22 confers different resistance responses against tobacco mosaic virus dependent on its expression level. Mol Plant 6, 971–974, doi:10.1093/mp/sss153 (2013).

60 Monino-Lopez, D. et al. Allelic variants of the NLR protein Rpi-chc1 differentially recognise members of the Phytophthora infestans PexRD12/31 effector superfamily through the leucine-rich repeat domain. Plant J, doi:10.1111/tpj.15284 (2021).

61 Castel, B. et al. Diverse NLR immune receptors activate defence via the RPW8-NLR NRG1. New Phytol 222, 966–980, doi:10.1111/nph.15659 (2019).

62 Sun, X. et al. Pathogen effector recognition-dependent association of NRG1 with EDS1 and SAG101 in TNL receptor immunity. bioRxiv, 2020.2012.2021.423810, doi:10.1101/2020.12.21.423810 (2020).

63 Lapin, D. et al. A Coevolved EDS1-SAG101-NRG1 Module Mediates Cell Death Signaling by TIR-Domain Immune Receptors. Plant Cell 31, 2430–2455, doi:10.1105/tpc.19.00118 (2019).

64 Gantner, J., Ordon, J., Kretschmer, C., Guerois, R. & Stuttmann, J. An EDS1-SAG101 Complex Is Essential for TNL-Mediated Immunity in Nicotiana benthamiana. Plant Cell 31, 2456–2474, doi:10.1105/tpc.19.00099 (2019).

65 Sun, X. et al. Pathogen effector recognition-dependent association of NRG1 with EDS1 and SAG101 in TNL receptor immunity. Nat Commun 12, 3335, doi:10.1038/s41467-021-23614-x (2021).

