## Supplemental tables, methods and materials for "Novel effector recognition capacity engineered into a paired NLR complex"

**This PDF file includes:**

**Tables S1 to S4**

**Material and methods**

**Table S1: effectors which target and degrade the host targets.**

| **Effector** | **Pathogen** | **Target proteins** | **Minimum interaction domain** | **Reference** |
| --- | --- | --- | --- | --- |
| SAP54 | *Phytoplasma* | MADS-domain transcription factor (MTF)  family, SEP3, AP1, SOC1 | Keratin like domain | ^1^ |
| SAP11 | *Phytoplasma* | TCP transcription factors | TCP domain | ^2^ |
| SAP05 | *Phytoplasma* | GATA/SBP transcription factors | GATA domain, SBP domain | ^3^ |
| HopX1 | *Pseudomonas syringae* | JAZ protein | JAZ592–163 (JAZ5 ZIM) | ^4^ |
| HopBB1 | *Pseudomonas amygdali* | TCP14, JAZ protein | TCP14(aa180-216), JAZ3 domain (206–302) | ^5^ |
| HaRxL44 | *Hyaloperonospora arabidopsidis* | MED19a |  | ^6^ |

**Table S2.** **The proteins sequences used in this study**

| Protein name | Sequence | Note |
| --- | --- | --- |
| [GS]_10_ | GSGSGSGSGSGSGSGSGSGS | Linker |
| [NAAIRS]_4_ | NAAIRS NAAIRS NAAIRS NAAIRS | Linker |
| [GS]_4_-[NAAIRS]_2-_-[GS]_4_ | GSGSGSGS NAAIRS NAAIRS GSGSGSGS | linker |
| GATA domain | SLLARRCANCDTTSTPLWRNGPRGPKSLCNACGIRFKKEERRTTAATGNTVVG | AtGATA18, AT3G50870 |
| SBP domain | CQIDGCELDLSSAKGYHRKHKVCEKHSKCPKVSVSGLERRFCQQCSRFHAVSEFDEKKRSCRKRLSHHNARRRK | putative squamosa-promoter binding protein 2, At1g27360 |
| keratin-like domain (KD) | KDRVSTKPVSEENMQHLKYEAANMMKKIEQLEASKRKLLGEGIGTCSIEELQQIEQQLEKSVKCIRARKTQVFKEQIEQLKQKEKALAAENEKLSEKWGS | keratin-like domain of AtSOC1, AT2G45660 |
| MED19A | MEPERLKFGGPRELCGAADLISQFKLVQHHEFFCKKSLPVSLSDSHYLHNVVGDTEIRKGEGMQLDQLIESISQSRETNIRIQPFDIDELQESFQLNDMTPVELPPAEKGAPTIPSKSKSESKDRDRKHKKHKDRDKDKDREHKKHKHKHKDRSKDKDKDKDRDRKKDKNGHHDSGDHSKKHHDKKRKHDGDEDLNDVQRHKKNKHKSSKLDEVGAIRVAG | AtMED19A, AT5G12230 |
| TCP2d | KDRHSKVLTSKGPRDRRVRLSVSTALQFYDLQDRLGYDQPSKAVEWLIKAAEDSISELP | TCP domain from TCP2, AT4G18390 |
| TCP18d | TDRHSKIKTAKGTRDRRMRLSLDVAKELFGLQDMLGFDKASKTVEWLLTQAKPEIIKIA | TCP domain from TCP18, AT3G18550 |
| JAZ1 | MSSSMECSEFVGSRRFTGKKPSFSQTCSRLSQYLKENGSFGDLSLGMACKPDVNGTLGNSRQPTTTMSLFPCEASNMDSMVQDVKPTNLFPRQPSFSSSSSSLPKEDVLKMTQTTRSVKPESQTAPLTIFYAGQVIVFNDFSAEKAKEVINLASKGTANSLAKNQTDIRSNIATIANQVPHPRKTTTQEPIQSSPTPLTELPIARRASLHRFLEKRKDRVTSKAPYQLCDPAKASSNPQTTGNMSWLGLAAEI | AtJAZ1, AT1G19180 |
| TCP14 | MQKPTSSILNVIMDGGDSVGGGGGDDHHRHLHHHHRPTFPFQLLGKHDPDDNHQQQPSPSSSSSLFSLHQHQQLSQSQPQSQSQKSQPQTTQKELLQTQEESAVVAAKKPPLKRASTKDRHTKVDGRGRRIRMPALCAARVFQLTRELGHKSDGETIEWLLQQAEPSVIAATGTGTIPANFTSLNISLRSSGSSMSLPSHFRSAASTFSPNNIFSPAMLQQQQQQQRGGGVGFHHPHLQGRAPTSSLFPGIDNFTPTTSFLNFHNPTKQEGDQDSEELNSEKKRRIQTTSDLHQQQQQHQHDQIGGYTLQSSNSGSTATAAAAQQIPGNFWMVAAAAAAGGGGGNNNQTGGLMTASIGTGGGGGEPVWTFPSINTAAAALYRSGVSGVPSGAVSSGLHFMNFAAPMAFLTGQQQLATTSNHEINEDSNNNEGGRSDGGGDHHNTQRHHHHQQQHHHNILSGLNQYGRQVSGDSQASGSLGGGDEEDQQD | AtTCP14, AT3G47620 |
| Unstructured domain | PRLRYQPLLRISQNCEAAILRASQTRLNTIGAYGSTVPRSQSFE | ^7^ |
| SAP05 | MAPNEEFVGDMRIVNVNLSNIDILKKHETFKKYFDFTLTGPRYNGNIAEFAMIWKIKNPPLNLLGVFFDDGTRDDEDDKYILEELKQIGNGAKNMYIFWQYEQK | Phytoplasma effecter |

**Table S3. Plasmids used in this study.**

| Plasmids | Vector | Purpose |
| --- | --- | --- |
| *35S:* RRS1-R-HF |  | WB, HR |
| *35S:* RRS1-R*^slh1^*-HF |  | WB, HR |
| *35S:* RRS1-R*^SH/AA^*-HF |  | WB, HR |
| *35S:*RPS4-HA |  | WB, HR |
| *35S:* RRS1-R*^slh1^*-V5 | pICSL86977OD | WB, HR |
| *pAT2*: RRS1-R*^slh1^*-V5 | pICH47742 | Transgenic Arabidopsis |
| *pAT2*: RRS1-R^SH/AA^-HF | pICH47751 | Transgenic Arabidopsis |
| *pAT2*: RRS1-R^SH/AA^-[GS]_10_-GATA HF | pICH47751 | Transgenic Arabidopsis |
| *35S:* RRS1-R-[GS]_10_-GATA-HF | pICSL86977OD | WB, HR |
| *35S:* RRS1-R-[GS]_10_-KD-HF | pICSL86977OD | HR |
| *35S:* RRS1-R-[NAAIRS]_4_-KD-HF | pICSL86977OD | HR |
| *35S:* RRS1-R-[GS]_4_-[NAAIRS]4_4_-[GS]_4_-KD | pICSL86977OD | HR |
| *35S:* RRS1-R^SH/AA^-[GS]_10_-TCP2d-HF | pICSL86977OD | WB, HR |
| *35S:* RRS1-R^SH/AA^-[GS]_10_-TCP18d-HF | pICSL86977OD | WB, HR |
| *35S:* RRS1-R^SH/AA^-[GS]_10-_GATA-HF | pICSL86977OD | WB, HR |
| *35S:* RRS1-R^SH/AA^-[GS]_10_-MED19a-HF | pICSL86977OD | WB, HR |
| *35S:* RRS1-R^SH/AA^-[NAAIR]_4_-GATA-HF | pICSL86977OD | WB, HR |
| *35S:* RRS1-R^SH/AA^-[NAAIR]_4_-SBP-HF | pICSL86977OD | HR |
| *35S:* RRS1-R^SH/AA^-[GS]_10-_SBP-HF | pICSL86977OD | HR |
| *35S:* RRS1-R^SH/AA^-[GS]_4_-[NAAIR]_2_-[GS]_4_-KD | pICSL86977OD | HR |
| *35S:* RRS1-R-[GS]_10_-TCP14 | pICSL86977OD | HR |
| *35S:* RRS1-R-[GS]_10_-JAZ1 | pICSL86977OD | HR |
| 35S: SAP05 | pICSL86977OD | WB, HR |

**Table S4. Primers used in this study.**

| Primer name | Sequence 5’-3’ |
| --- | --- |
| *GATA-F* | *TCTCTTCTCGCTAGACGCTGTG* |
| *GATA-R* | *CCTCCGACGACGGTGTTTCCTGTAG* |
| *KD-F* | *AAGGATCGAGTCAGCACCAAACCGGTTTC* |
| *KD-R* | *TCAAGATCCCCACTTTTCAGA* |
| *TCP2d-F* | AAAGATCGACACAGCAAAGTCTT |
| *TCP2d-R* | AGGAAGCTCAGAGATTGAATCT |
| *TCP18d-F* | ACGGACAGGCACAGCAAGATCAAA |
| *TCP18d-R* | CGCGATCTTTATGATCTCAGGT |
| *MED19a-F* | ATGGAGCCTGAACGTTTAAAATTTG |
| *MED19a-R* | GCCAGCAACCCTTATTGCACCCAC |
| *SBP-F* | TGCCAAATTGATGGCTGTGAA |
| *SBP-R* | CCTTACGACGCCTCGCATTATGATG |
| *TCP14-F* | ATGCAAAAGCCAACATCAAGTATC |
| *TCP14-R* | ATCTTGCTGATCCTCCTC |
| *JAZ1-F* | ATGTCGAGTTCTATGGAATGTTC |
| *JAZ1-R* | TCATATTTCAGCTGCTAAACCG |
| *SAP05-F* | ATGGCCCCGAATGAAGAGTTT |
| *SAP05-R* | TTACTTTTGCTCATATTGCCAGAA |

**NOTE:** the listed sequences are the gene-specific sequences, the enzyme recognition sites, generates complementary overhangs are not included.

**Material and methods**

**Plasmid construction**

All plasmids generated in this study are listed in the SI Appendix Table S3. All the protein sequence are listed in the SI Appendix Table1. And all the primers used in this study are listed in SI Appendix Table S2. The templates are from the published study. Phusion® High-Fidelity DNA Polymerase (Cat No. M0530S) from NEB was used for PCR amplification. BsaI (Cat No. ER0291) and BpiI (Cat No. ER1011) used in Goldengate cloning were purchased from Thermo Fisher Scientific. T4 DNA ligase (Cat. M0202S) and Bovine Serum Albumin (BSA) (Cat. B9000S) were purchased from NEB. Escherichia coli strains DH5α and Agrobacterium tumefaciens strain GV3101 or Agl1 were used for transformation and were grown on the Luria-Bertani (LB) medium plate at 37 °C and 28 °C, respectively.

**Plant growth condition**

*Nicotiana tabacum* cv. “Petite Gerard” and *N. benthamiana* were grown in growth chamber in soil at 24 °C, 55% relative humidity with a 16/8-h light/dark photoperiod. *Arabidopsis thaliana* was grown in growth chamber in soil at 21 °C, 70% relative humidity with a10/14-h light/dark photoperiod.

**Leaf Infiltration and HR assay**

HR assay were carried out on 4 to 5-week-old *N. tabacum* lamina sections between veins by agroinfiltration. Protein assay were performed in 4 to 5-week-old *N. benthamiana* plants. Agrobacterium tumefaciens strains were suspended and mixed in infiltration buffer (10 mM MgCl2, 10 mM 2-(N-morpholino) ethanesulfonic acid [MES], pH 5.6) with each at an OD_600_=0.5 (agrobacteria strain with SAP05 was infiltrated at OD_600_ = 1), and hand-infiltrated with a 1 mL needless syringe. Three to four days after infiltration, HR was scored based on the HR_Index defined in Fig. 1f.

**Phytoplasma infection**

Phytoplasma infection assay with Arabidopsis is performed as reported ^3^. To be brief, one expanded leaf from 4-week-old Arabidopsis was exposed to 2 or 3 leafhoppers (*M. quadrilineatus* colonies carrying the AY-WB phytoplasma) in a clip cage for 2-3 days. The insects were removed from plants after 3 or 4 days. Disease symptom progression was monitored starting from 3 weeks after insect exposure.

**Protein assays**

Protein was extracted from *N.benthamiana* leaves 48-72 hours post infiltration (hpi). For total protein detection, plant leaves were harvest and ground in liquid nitrogen, and boiled in SDS loading buffer with 10 mM 1,4-dithiothreitol (DTT). For co-immunoprecipitation assay, leaves were harvested 48 hpi and ground in liquid nitrogen, and extracted in GTN buffer (10% glycerol, 75 mM Tris·HCl, pH 7.5, 150 mM NaCl, 5 mM 1,4-dithiothreitol (DTT), 1x cOmplete™ protease inhibitor cocktail [Roche] and 0.5% Nonidet™ P-40 [Sigma-Aldrich]). Samples were then centrifuged at 4,000 rpm, 4 ⁰C and the supernatant was used for immunoprecipitation and for input. Immunoprecipitation were performed at 4 ⁰C, and the supernatant was incubated with anti-FLAG M2 affinity Gel (A2220 sigma-Aldrich) for 1 hour. Beads were washed 3 times with GTN buffer after incubation and then boiled in 2x SDS-PAGE loading buffer. ANTI-FLAG® M2-Peroxidase (HRP) (Sigma-Aldrich) antibody, V5-HRP antibody (Invitrogen) and anti-SAP05 antibody (from Saskia Hogenhout lab) ^3^ were used for immunoblotting.

**Statistics analysis**

All the statistical tests used are two-tailed. The original data, results of all the statistical analysis and script can be found in DataSet_01.

For HR assay, the one-way analysis of variance (ANOVA), followed by Turkey HSD test is used to determine whether there are any statistically significant differences. Spreadsheet 4, 5 and 6 are the original scores for the corresponding HR assays. The F values and degrees of freedom for the ANOVAs analysis of HR assays are listed in spreadsheet7.

For phytoplasma infection assay, Student’s *t* test is used to test if the proportion of plants with severe symptom are different between each group with *p* value labelled on the plot. The error bar shows the 95% confidence interval. In the DataSet_01, spreadsheet1 to 3 are related to Fig. 3d. The *p* value, 95% confidence interval, t-value and degree of freedom are listed in spreadsheet3.

**Reference**

1	MacLean, A. M. *et al.* Phytoplasma effector SAP54 hijacks plant reproduction by degrading MADS-box proteins and promotes insect colonization in a RAD23-dependent manner. *PLoS Biol* **12**, e1001835, doi:10.1371/journal.pbio.1001835 (2014).

2	Pecher, P. *et al.* Phytoplasma SAP11 effector destabilization of TCP transcription factors differentially impact development and defence of Arabidopsis versus maize. *PLoS Pathog* **15**, e1008035, doi:10.1371/journal.ppat.1008035 (2019).

3	Huang, W. *et al.* Parasite co-opts a ubiquitin receptor to induce a plethora of developmental changes. *bioRxiv*, 2021.2002.2015.430920, doi:10.1101/2021.02.15.430920 (2021).

4	Gimenez-Ibanez, S. *et al.* The bacterial effector HopX1 targets JAZ transcriptional repressors to activate jasmonate signaling and promote infection in Arabidopsis. *PLoS Biol* **12**, e1001792, doi:10.1371/journal.pbio.1001792 (2014).

5	Yang, L. *et al.* Pseudomonas syringae Type III Effector HopBB1 Promotes Host Transcriptional Repressor Degradation to Regulate Phytohormone Responses and Virulence. *Cell Host Microbe* **21**, 156-168, doi:10.1016/j.chom.2017.01.003 (2017).

6	Caillaud, M. C. *et al.* A downy mildew effector attenuates salicylic acid-triggered immunity in Arabidopsis by interacting with the host mediator complex. *PLoS Biol* **11**, e1001732, doi:10.1371/journal.pbio.1001732 (2013).

7	Fishbain, S., Prakash, S., Herrig, A., Elsasser, S. & Matouschek, A. Rad23 escapes degradation because it lacks a proteasome initiation region. *Nat Commun* **2**, 192, doi:10.1038/ncomms1194 (2011).
